# Bayesian ^13^C-metabolic flux analysis of parallel tracer experiments in granulocytes: A directional shift within the non-oxidative pentose phosphate pathway supports phagocytosis

**DOI:** 10.1101/2023.11.01.565126

**Authors:** Melanie Hogg, Eva-Maria Wolfschmitt, Ulrich Wachter, Fabian Zink, Peter Radermacher, Josef Albert Vogt

**Affiliations:** Institute for Anesthesiological Pathophysiology and Process Engineering, Ulm University Medical Center, Ulm, Germany

**Keywords:** Bayesian modeling, gas chromatography-mass spectrometry, glucose metabolism, mass isotopomer distribution analysis, sugar phosphates

## Abstract

The pentose phosphate pathway (PPP) plays a key role in the cellular regulation of immune cell function; however, little is known about the interplay of metabolic adjustments in granulocytes, especially regarding the non-oxidative PPP. For the determination of metabolic mechanisms within glucose metabolism, we propose a novel Bayesian ^13^C-Metabolic flux analysis based on *ex-vivo* parallel tracer experiments with [1,2-^13^C]glucose, [U-^13^C]glucose, and [4,5,6-^13^C]glucose and gas chromatography-mass spectrometry labeling measurements of metabolic fragments including sugar phosphates. With this approach we obtained precise flux distributions and their joint confidence regions, which showed that phagocytic stimulation reversed the direction of non-oxidative PPP net fluxes from ribose-5-phosphate biosynthesis towards glycolytic pathways. This process was closely associated with the up-regulation of the oxidative PPP to promote the oxidative burst. The estimated fluxes showed strong pairwise inter-relations forming a single line in several cases. This behavior could be explained with a three-dimensional permissible space derived from stoichiometric-flux-constraint analysis and enabled a principal component analysis detecting only three distinct axes of coordinated flux changes that were sufficient to explain all flux observations.

## 1 Introduction

Granulocytes/neutrophils are the most abundant immune cells in circulation. They play a key role in the innate immune system and are the first responders of defense against pathogens (Kumar & Dikshit, 2019). On the site of infection, neutrophils ensnare microbes to kill them by oxidative burst or neutrophil extracellular traps (NET) formation (Toller-Kawahisa & O’Neill, 2022). To fulfill these effector functions upon stimulation, neutrophils quickly (as early as 10 min) adapt their metabolic program from glycolysis with minor oxidative PPP activity to pentose phosphate recycling to maximize oxidative PPP activity (Britt et al, 2022). In neutrophils, the oxidative PPP is the main source of NADPH for reactive oxygen species (ROS) production *via* the NADPH oxidase (Paclet et al, 2022; TeSlaa et al, 2023). Therefore, an increased oxidative PPP activity after stimulation can be interpreted as a key indicator for granulocyte activation. This process is usually accompanied by a reduction of main glycolytic pathways due to competition for the substrate glucose-6-phosphate (G6P). The same effect can be caused by glycolytic mediators like TIGAR (TP53-Induced glycolysis and apoptosis regulator), which decreased ROS levels by promoting PPP activity in human lymphocytes (Simon-Molas et al, 2021; Stanton, 2012). Substrate availability further affects the direction of the non-oxidative PPP: In a series of reversible reactions, its enzymes transaldolase and transketolase convert pentose phosphates originating from the oxidative PPP into glycolytic intermediates including fructose-6-phosphate (F6P) and glyceraldehyde-3-phosphate (GAP). Non-oxidative PPP reactions in this direction complete the oxidative PPP and have been most widely studied, but reactions in the opposite direction are equally likely and can even occur simultaneously (Li et al, 2019). This flexibility could transform the non-oxidative PPP into a key element of anabolic and catabolic glucose metabolism (Qi Liu et al, 2022). For example, Jeon *et al*. highlighted that “a detailed study on the role of the non-oxidative branch of PPP in neutrophils is still needed” (Jeon et al, 2020).

There are additional interplaying metabolic mechanisms in granulocytes that can drastically complicate the investigation of glycolytic pathways and PPP, like gluconeogenesis and the glycogen synthesis/degradation cycle (Sadiku et al, 2021). The reversibility of reactions and the vast interactions of the network call for an elaborate method to capture the entire metabolic program. To tackle this challenge, we have employed a gas chromatography-mass spectrometry (GC-MS)-based ^13^C metabolic flux analysis (^13^C-MFA). ^13^C-MFA is a powerful tool for flux quantification including reversible and irreversible reactions. However, increased computational and technical challenges are associated with the reversibility of fluxes and the strong interactions between PPP and glycolysis as the complexity of the network increases (Zamboni et al, 2009). The accuracy and precision of glucose-centered ^13^C-MFA relies on ^13^C labeling measurements of intracellular metabolites like sugar phosphates, whose detection is a challenging task due to low intracellular levels and the occurrence of multiple isomers. In previous studies, the labeling data from ^13^C tracer experiments were most commonly determined by liquid chromatography-mass spectrometry (LC-MS and LC-MS/MS) and recently by gas chromatography-negative chemical ionization-mass spectrometry (GC-NCI-MS) (Hanke et al, 2013; Britt et al, 2022; Haschemi et al, 2012; Sadiku et al, 2021; Rühl et al, 2012; Okahashi et al, 2019). In addition to LC-MS being costly and struggling with baseline separation of hexose phosphates, both LC-MS and GC-NCI-MS techniques only provide labeling information of the entire carbon skeleton, resulting in the loss of valuable positional information for flux resolution, e.g. location of label concentration. In contrast, GC-MS with electron ionization (EI) provides multiple fragments with the potential to increase information for ^13^C-MFA (Wiechert, 2001). However, the extensive fragmentation may lead to low sensitivity for certain fragment ions, especially for high *m/z* values like fragment ions containing the entire carbon skeleton of sugar phosphates. The combination of low abundance of sugar phosphates and EI-fragmentation raises the question whether the provided measurement precision is sufficient for reliable flux determination. Therefore, strategies to combat low sensitivity and the selection of optimal ^13^C-labeled glucose tracers to provide essential labeling information are critical components for quantification of metabolic fluxes. As the gold standard in ^13^C-MFA, different tracers can be studied in parallel incubations under comparable conditions, which allows a comprehensive validation and offers the potential to increase flux accuracy and precision (Crown et al, 2015; Antoniewicz, 2013; Antoniewicz, 2015). However, limitations arise due to the increase in required sample material, which is particularly challenging in *ex-vivo* ^13^C-MFA approaches. We have chosen [1,2-^13^C]glucose, [4,5,6-^13^C]glucose, and [U-^13^C]glucose for our set up to simultaneously capture the oxidative and non-oxidative PPP, gluconeogenesis, and tracer dilution. The latter was especially important to detect sources of unlabeled carbon into the PPP; either frequently considered sources like the glycogen synthesis/degradation cycle, or more theoretical inputs into triose, pentose or sedoheptulose (Sadiku et al, 2021; Haschemi et al, 2012; Nagy & Haschemi, 2013) The combination of GC-EI-MS with parallel tracer experiments enabled detection of various fragments including sugar phosphates that are essential for reliable flux determination and sufficient flux resolution of glycolysis and the PPP. When adapting a MFA to a Bayesian analysis, we obtain not only a single, fixed flux value but flux distributions and their common, potentially nonlinear confidence regions. This enables easy plotting of pairwise couplings; and for a single sample we can infer the extent to which the determination of two fluxes is coupled given the underlying data set. For differences observed when comparing two or more samples or groups, Bayesian analysis facilitates the assessment whether these differences can be explained by determination uncertainty alone or whether there is an underlying biological effect.

We therefore used this method to unravel and quantify the interplay of key processes of glucose metabolism by revealing coordinated changes with a principal component analysis (PCA). Decisive pathways within the fluxes were deduced from the “Stoichiometric Constraint Matrix” provided by our flux balances in the Bayesian ^13^C-MFA, where input equals output for each metabolite pool. As input for the PCA, we utilized individual ^13^C-MFA data from granulocytes (n = 28) of different degrees and methods of stimulation (untreated, *E.coli* bioparticle-stimulated, PMA-treated, PMI+DPI-treated) and investigated whether the metabolic plasticity is limited to so few processes that changes in the individual fluxes can be detected with the given measurement accuracy.

## 2 Materials and methods

### 2.1 Materials

RPMI 1640 powder (without glutamine, glucose, NaHCO_3_) was produced by Genaxxon bioscience (Ulm, Germany) and the derivatization reagents N,O-bis(trimethylsilyl)- trifluoroacetamide (BSTFA) were purchased from abcr (Karlsruhe, Germany). Pig serum was obtained from Bio-Rad Laboratories (Hercules, CA, USA) and Pancoll human (density of 1.119 g/mL and density of 1.077 g/mL) were purchased from PAN Biotech (Aidenbach, Germany). The pHrodo™ Green *E. coli* BioParticles™ phagocytosis kit for flow cytometry and phosphate buffer saline (PBS, without Ca^2+^, Mg^2+^) were produced by Thermo Fisher Scientific (Waltham, MA, USA). We obtained Ampuwa (aqua ad iniectabilia) and sterile 0.9% NaCl solution from Fresenius Kabi (Bad Homburg, Germany). [1,2-^13^C]glucose (99 atom% ^13^C), [4,5,6-^13^C]glucose (99.5%), [U-^13^C]glucose (99%), [U-^13^C]glucose-D-6-phosphate disodium salt hydrate (99%) were purchased from Cambridge Isotope Laboratories (Andover, MA, USA). All other chemicals and standard substances were purchased from Sigma-Aldrich (St. Louis, MO, USA).

### 2.2 Preparation of RPMI tracer medium

Modified RPMI-1640 medium was prepared as follows: one aliquot of RPMI 1640 powder suitable for 1 L RPMI was completely dissolved in 900 mL Ampuwa water. 4.766 g HEPES corresponding to a final concentration of 20 mM was added before adjusting the pH to 7.5 with 1 N NaOH. The RPMI stock solution was adjusted to 1 L with Ampuwa water, sterilized using a 100 µm sterile filter and stored at 4°C until isotopic tracer experiment. For experiments involving tracer mixtures, stock RPMI medium (pH: 7.5) was supplemented with 0.1 mg/mL sodium bicarbonate, 0.6 mg/mL glutamine, 0.9 mg/mL glucose and 0.9 mg/mL of the corresponding isotopic tracer ([1,2-^13^C]glucose, [4,5,6-^13^C]glucose, [U-^13^C]glucose) or unlabeled glucose for phagocytosis assay.

### 2.3 Isolation of granulocytes from whole blood

Blood collection of experimental animals was performed after obtaining the approval by the University of Ulm Animal Care Committee and the Federal Authorities for Animal Research (Regierungspräsidium Tübingen; Reg.-Nr. 1559, approval October 29, 2021) and in compliance with the National Institute of Health Guidelines on the Use of Laboratory Animals and the European Union “Directive 2010/63/EU on the protection of animals used for scientific purposes”. Arterial blood from adult, human-sized German landrace swine was collected immediately after anesthesia and instrumentation as described in detail by Münz *et al*. (Münz et al, 2023). Directly after sampling, granulocytes were isolated from the lithium-heparin blood using Ficoll density centrifugation. Briefly, blood was diluted with an equal volume of PBS, layered on two-density gradient solution (density: 1.077 g/mL, 1.119 g/mL (9:8, v/v)), and centrifuged at 800 g (20 min without break at room temperature (RT)). The upper layer containing Ficoll and plasma was discarded, while the lower fraction containing red blood cells and granulocytes was transferred to a new tube. Osmotic lysis was performed by adding ice cold water, incubating for 2 min, and then stopping the reaction by addition of 10× PBS. This step was performed three times to purify the granulocytes before counting cells in a Neubauer counting chamber. All experiments comprised 10 biological replicates. Phagocytosis assay and isotopic tracer experiments were performed within 2-3 h after blood collection.

### 2.4 Phagocytosis assay

For stimulation, 5 × 10^6^ purified granulocytes were washed with 1 mL of ice cold RPMI 1640 medium. Cells were resuspended in 200 µL of ice cold RPMI containing 10% pig serum. Next, 50 µL of *E.coli* BioParticles (1 mg/mL) was added to the cell suspension und incubated for 20 min at 37°C. After stimulation, 1 mL ice cold phagocytosis washing buffer was added and the cells were harvested by centrifugation (5 min, 400 g, 4°C).

### 2.5 Isotopic tracer experiments

For parallel isotopic labeling experiments, 5 × 10^6^ purified granulocytes with and without prior phagocytosis assay were resuspended in 1 mL RPMI medium spiked with isotopic tracer containing [1,2-^13^C]glucose, [4,5,6-^13^C]glucose, or [U-^13^C]glucose (labeled/unlabeled, 1:1, *w/w*), respectively. The incubation was performed in a closed 2 mL tube at 37°C under constant vertical rotation at 5 rpm (Trayster Digital, IKA, Staufen, Germany) for 2 h. Recently, an *ex-vivo* ^13^C analysis of neutrophils showed that a steady state of the PPP/glycolytic fluxes was achieved between 10 - 30 min (Britt et al, 2022). Another study of Kuehne *et al*. demonstrated that the steady state of G6P ^13^C labeling was observed within 10 min when investigating the PPP in human skin (Kuehne et al, 2015). Based on these previous findings and the short life cycle (< 1 day) of granulocytes, we assumed an isotopic steady state, constant fluxes, and stable labeling patterns within 2 h. After incubation, cells were washed once with 0.9% NaCl solution and subsequently stored at −80°C after removal of all liquid. Samples were stored for about 2 weeks until analysis.

### 2.6 Extraction of intracellular metabolites

The frozen pellet was resuspended in 100 µL ice-cold Ampuwa water, vortexed vigorously and sonicated in an ice bath for 10 min. Next, 400 µL of cold acetonitrile/ methanol (1:1, *v*/*v*, −20°C) was added to the suspension. After extraction for 10 min using an ultrasonic cleaning bath, samples were centrifuged at 14 000 g and 4 °C for 5 min. The supernatants from duplicates of the isotopic tracer experiments were pooled in a 1.5 mL vial. The combined extract was dried at 45°C using the SpeedVac evaporator (Savant SPD2010 SpeedVac concentrator, Thermo Scientific, Waltham, MA, USA). The residue was kept overnight at −20°C until derivatization and GC-MS analysis.

### 2.7 Derivatization and GC-MS analysis of intracellular metabolites

The cell extracts and reference standard mix containing individual concentrations of various sugar phosphates (G6P, F6P, G1P, M6P, X5P, Ru5P, R5P, DHAP, 3PG, G3P), glucose, and lactate were analyzed as ethyloxime-trimethylsilyl derivatives (EtOx-TMS). For this approach, the dried cell extracts and five staggered levels of reference standard mix (0.006 to 6 µg) were dissolved in 50 µL solution of ethoxyamine hydrochloride in pyridine (2%, *w*/*v*). After sonication for 10 min and incubation at 60°C for 60 min, the samples were evaporated with a gentle stream of nitrogen at 45°C. The dried residue was dissolved in 30 µL acetonitrile and 30 µL BSTFA in the ultrasonic cleaning bath (15 min). For derivatization, the reaction mixture was heated to 60°C for 45 min. Finally, the sample was centrifuged for 5 min at 14000 g before transferring the clear liquid into a GC vial for GC-MS analysis.

All samples were analyzed on a 7890A/5977B GC-MS system (Agilent, Waldbronn, Germany) equipped with a 30-m OPTIMA^®^ 1301-MS column (6% cyanopropylphenyl, 94% dimethylpolysiloxane, 0.25 mm internal diameter, 0.25 µm film thickness; Macherey-Nagel, Düren, Germany). Injections of 1 µL aliquots of sample solution were performed in splitless or split mode depending on the peak intensities (n = 2, repetition measurements). The mass spectrometer was operated in EI mode at 70 eV. For the determination of the mass isotopomer distribution, the MS was operated in selected ion monitoring (SIM) mode (SIM parameters and GC-MS settings are provided in Supplementary file: research_data.xlsx). Integration was performed with our in-house program: A seven parameter model was used to describe the peak shape, with both legs of the peak approaching zero. For a given metabolite fragment, mean and standard deviation of all peak parameters were determined from measurements of all relevant masses of at least 5 reference samples. The peak model was used to integrate a measured elution curve for a given *m/z* value of a study sample with the specification that the peak parameters remain within the range of mean and standard deviation of the reference samples. The same time window for integration was used for reference and study samples. Since the model elution curve tends to zero at both ends, a straight line was added to it as a baseline to better track the measured curve. The carbon mass distribution (CMD) was obtained after correction for naturally occurring isotopes like nitrogen, oxygen, hydrogen and silicon with a correction matrix approach (van Winden et al, 2002; Wolfschmitt et al, 2023). The CMDs were corrected for natural abundance of ^13^C for model free evaluation, but not for the modeling-based MFA, which had natural ^13^C included in its ^13^C label definition.

### 2.8 Metabolic network model and metabolic flux analysis

#### 2.8.1 Glossary

- C1, C2, …C6: Designation of the position for the carbons of a metabolite.
- *c*_1_, *c*_2_, … *c*_*Nc*_: Isolated labeling on the carbons of a fragment metabolite.
- CMD: Normalized carbon mass distribution of a fragment with the elements *r*_*i*_, where *i* denotes the mass offset or number of simultaneously labeled carbons.
- MID: Normalized mass isotopomer distribution of a fragment without correction for naturally occurring isotopes of its elements (e.g., H, C, N, O, and Si). Values are expressed as mol%.
- *N*: Number of carbon labeling positions of a fragment metabolite.

#### 2.8.2 ^13^C-positional labeling approach

EI-MS provides the mass distributions across different fragments of the carbon skeleton. The following section shows how to derive positional labeling from the different fragments of an analyte.

The total ^13^C enrichment (*T*) of an ion fragment across the carbons *c*_*x*_ to *c*_*y*_ is denoted as

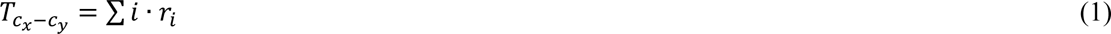

where *r*_*i*_ refers to an element of the CMD of the fragment of interest. Alternatively, the total ^13^C enrichment can be calculated from the sum of the isolated carbon labeling, i.e.

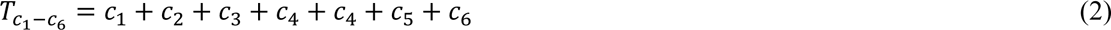

When observing the fractional ^13^C enrichment (*F*),

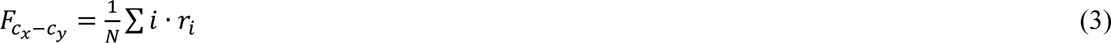

refers to the average carbon labeling of a fragment. In the situation when a molecular ion or larger fragment (e.g., C1-C6) breaks down into two smaller fragments (e.g., C1-C2 and C3-C6) we obtain:

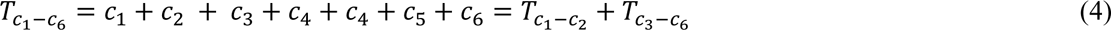

from the equations above. This equation can be arranged to

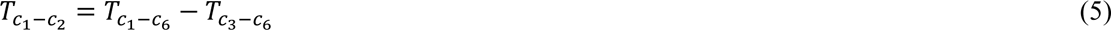

Thus, if a smaller fragment is cleaved from a larger one and both the initial and one of the cleaved fragments are measurable, then the difference of their total enrichments can be used to estimate the enrichment of the second fragment being formed (Lima et al, 2021).

#### 2.8.3 Accurate assessment of the selected fragments for ^13^C-MFA

GC-MS measurement accuracy was the main criterium of fragment selection for ^13^C-MFA. For a given fragment, the peak area of each mass isotopomer was normalized so that all signal areas from M+0 to M+(n+1) summed up to 1 (100 mol%):

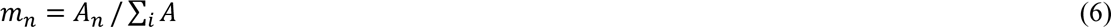

with *n*: number of carbons

*m*_*n*_: fractional abundance of each isotopomer of a fragment (M+0 to M+(n+1))

*A*_*n*_: peak area of an individual mass isotopomer

∑_*i*_*A*: sum of all signal areas

The corresponding theoretical MID were calculated with the isotope distribution calculator from Agilent (MassHunter Workstation Data Analysis Core, Version 8.0.8208.0, Agilent Technologies). The measured MID of each *m/z* of the fragments was compared with its theoretical value. Accuracy was assessed following Zamboni’s approach (Zamboni et al, 2009). An absolute error (difference) of up to 1.5 mol% was deemed acceptable for ^13^C-MFA, whereas an absolute error up to 0.8 mol% was optimal. The results for each analyzed fragment are shown in the Supplementary file: research_data.xlsx).

#### 2.8.4 Bayesian modeling for ^13^C-MFA

We established a Bayesian implementation of a combined model for glycolytic and PPP fluxes using the rstan package v2.21.2 (R interface to Stan) (Stan Development Team, 2020; Carpenter et al, 2017). It utilizes an adaptive and efficient Hamiltonian Monte Carlo sampler and provides several diagnostics that allow to check whether the inference is reliable. Its algorithm draws random samples of unknown parameters, such as flow rates. In this study, it subsequently computed CMDs from sampled fluxes by applying a user-defined model based on the elementary metabolite units (EMU) concept (Antoniewicz et al, 2007). If the EMU-calculated CMDs were comparable to the corresponding GC-MS measurements, the sample was collected in a Markov chain Monte Carlo (MCMC) sampling chain (Hastings, 1970); otherwise, it was disregarded.

The resulting sampling chain contains thousands of samples of the “true” posterior distribution, from which one can derive distributions of all estimated fluxes. The probability distribution “faithfully reports the full uncertainty due to experimental error, and any potential model data incompatibilities” (preprint: Backman et al, 2023) and relies on “all collected information about an unknown model parameter” (Theorell et al, 2017). We used a Dirichlet distribution to calculate the probability that a measured CMD can be explained by a calculated CMD. This is based on the following consideration: Molecules of a fragment are in a reservoir from which NC (number of counts) samples are drawn. If, for a given fixed NC, more counts are randomly drawn for one mass than expected, fewer will remain for the other masses. For normalization, the counts for a specific mass are divided by the total counts of all masses of the fragment. This inevitably results in a non-negligible correlation between the individual distribution elements. A Dirichlet distribution captures the resulting covariance. The variance (**σ**^2^) of a distribution element with abundance *M*_*i*_ is denoted as:

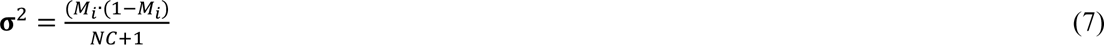

with *NC*: as the number of counts, considered in the precision parameter of a Dirichlet distribution.

The use of the Dirichlet distribution has the desired property that the measurement error for a fragment can be fixed with the single quantity NC. Moreover, the relative variance of a distribution element increases with decreasing abundance; therefore, there is a limit at which the variance becomes equal to the mean. However, the Dirichlet distribution only captures normalization-induced correlations, but for the mentioned urn scenario, a correlation between the distribution elements that already exists before sampling is not taken into account. Thus, the Dirichlet approach is only an approximate minimal model for error determination.

The estimated likelihood is maximal if the following two conditions are met: first, the difference between predicted and measured distributions should be minimal. Second, the theoretical standard deviations of the individual distribution elements should on average equal the difference between the elements of the theoretical and calculated CMD values; the precision for each fragment has to be set accordingly. More details on the adaption of Bayesian modeling for ^13^C-MFA are available in Supplementary files (Bayes_implementation.pdf, and EMU_level.xlsx).

#### 2.8.5 Stoichiometric flux restraint analysis of the metabolic network

For the MCMC sampling, flux samples were drawn from a reservoir of possible fluxes. This reservoir was defined by the requirements that i) fluxes must not become negative and ii) for each metabolite pool, the sum of incoming fluxes must equal the sum of outgoing fluxes. Two Bayesian ^13^C-MFA approaches from Theorell *et al*. and Borah Slater *et al*. have combined all flux constraints in a polytope from which a valid flux set was extracted (Theorell et al, 2017; Borah Slater et al, 2023). While being numerically expensive, the corresponding algorithms ensured that the entire valid domain was covered (Theorell et al, 2022; Zhang & Gao, 2003). However, the understanding of the range in which the fluxes can move was lost with the resulting "black box". We believe that this understanding would be critical to the interpretation of observed flux values and therefore specify the possible flux ranges in detail.

Flow rates were estimated based on the network shown in Figure 1. For this network, one can establish the following flux balance:

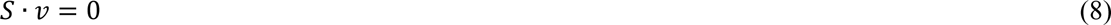

**Figure 1.**
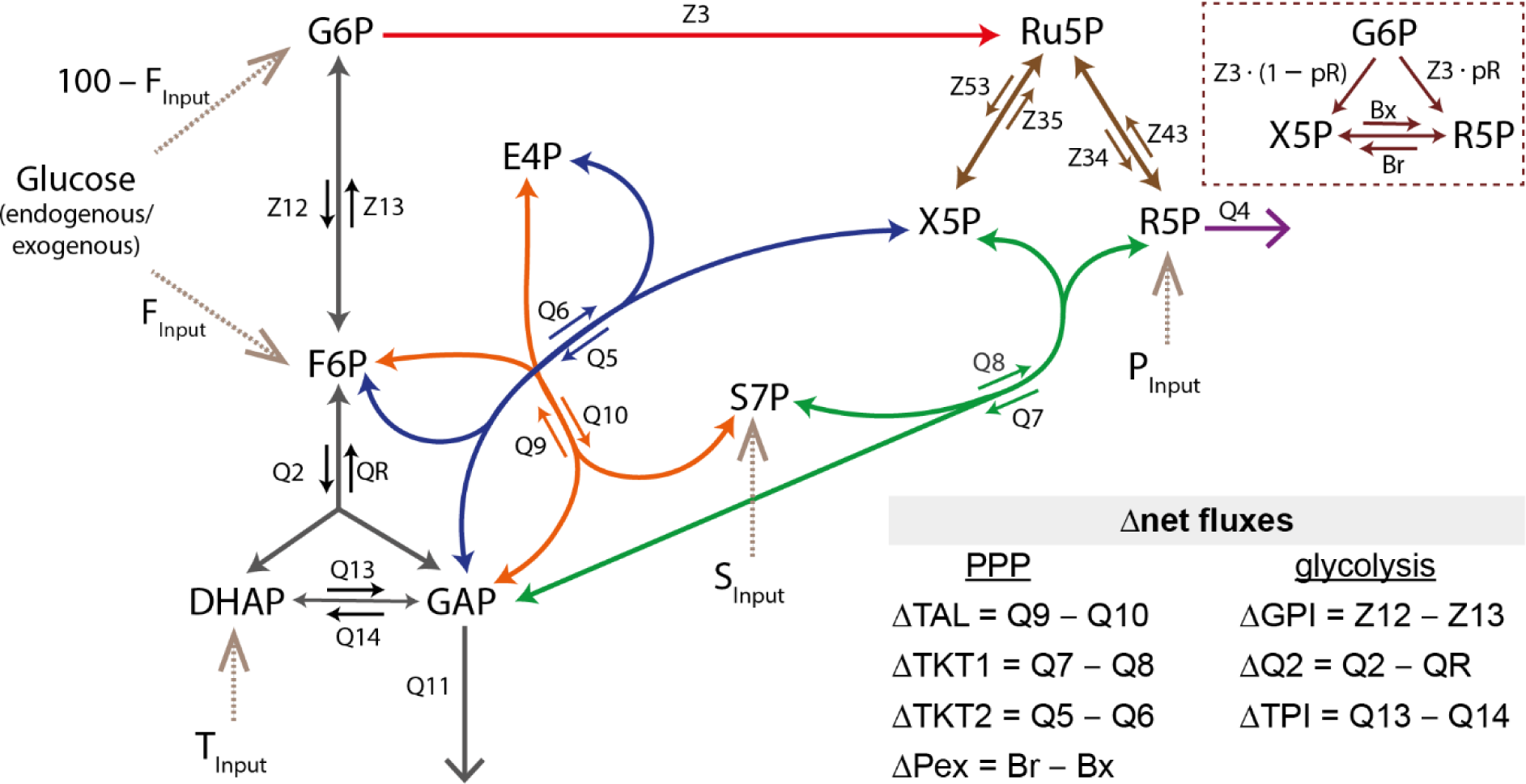
Metabolic network of the upper glucose metabolism for ^13^C-MFA. Fluxes were normalized to a glucose uptake rate of 100 into the combined F6P and G6P pool. Top right side: reduced system of the exchange between the pentose phosphate pool. Color code for flows: red: oxidative PPP, brown: pentose phosphate exchange, purple: R5P loss, green/blue: TKT1/2 reactions, orange: TAL reaction, black: glycolysis. The dashed lines indicate an input into the system, with P_Input_, S_Input_, and T_Input_ being unlabeled carbon sources. Abbr.: TAL = transaldolase, TKT = transketolase, P_ex_ = Pentose exchange, GPI = glucose-phosphate isomerase, TPI = triose-phosphate isomerase, metabolite abbr. see Supplementary file: Bayes_implementation.pdf

where *S* is a stoichiometry matrix with the number of rows equaling the number of pools and the number of columns equaling the number of fluxes. *v* contains the different fluxes. The stoichiometric equation can be divided into two components:

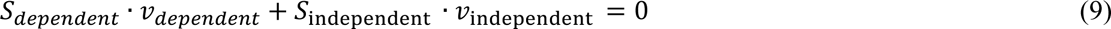

The number and selection of dependent fluxes must be chosen so that *S*_*dependent*_is a square matrix that is invertible. Thus, the equation above can be solved for the dependent fluxes:

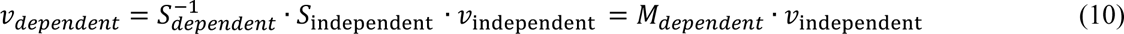

We derived a stoichiometry-restraint matrix (equation 11) which considered all irreversible and net fluxes of reversible reactions in our metabolic model (Figure 1). In the following, we used the symbol Δ for net fluxes of reversible reactions with the individual definitions presented in Figure 1. Dependent fluxes could be calculated from independent fluxes, which reduced the number of parameters and the permissible range of fluxes and therefore resulted in improved MCMC sampling. We have chosen ΔQ2, and ΔTAL as independent fluxes because these fluxes were candidates for control over the metabolic network. A third candidate for accessible control, the oxidative PPP (Z3), could not be defined as independent flux as this would lead to a non-invertible matrix in equation (10). Thus, we selected ΔQ2, ΔTAL, and all input fluxes as independent fluxes.

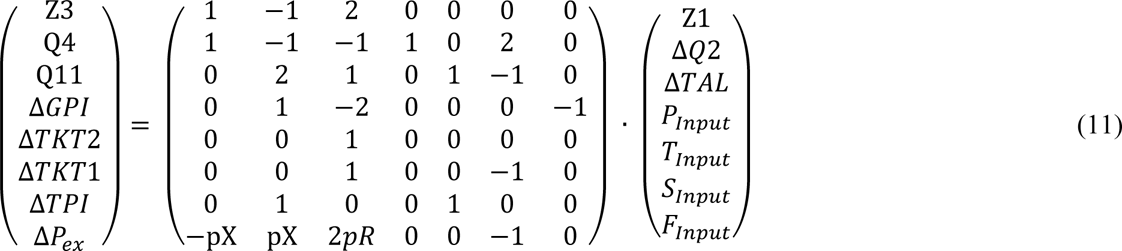

The choice of independent fluxes concluded that within the dependent fluxes only the oxidative PPP (Z3), the R5P loss (Q4), and the triose output (Q11) must be greater than or equal to zero. The resulting set of three equations derived from the first three rows of equation (11) could thus be considered the permissible space of the constraints. These equations depended on a maximum of 6 independent fluxes when various inputs were added, corresponding to the vector on the right side of equation (11) with Z1 being fixed at Z1 = 100. With independent fluxes drawn from the permissible space, the remaining 5 rows of equation (11) could be used to calculate the net fluxes. In a final step, net fluxes must be supplemented with exchange fluxes so that forward and backward fluxes were always greater than or equal to zero. The dimension of the restrain matrix generally equals the number of metabolite pools in the system (8 in the present case). By restricting the system to unidirectional and net fluxes we could reduce it to three dimensions, which allowed for efficient sampling. Details were given in Supplementary file: Bayes_implementation.pdf.

In the scenario of all input fluxes being zero, we speak of a “closed glycolysis/PPP system”. For this particular case, ΔTAL and ΔQ2 were the only independently variable fluxes, resulting in only two degrees of freedom.

#### 2.8.6 Evaluation of the Bayesian ^13^C-MFA

To test the accuracy of our ^13^C-MFA routine we evaluated the Bayesian model against a widely used INCA-software, which was applied by Britt *et al*. (Britt et al, 2022). To compare both approaches we used the Britt *et al*. ^13^C labeling data (3PG (C1-C3), DHAP(C1-C3), F-1,6-BP (C1-C6), 6PG (C1-C6), R5P (C1-C5), Ru5P(C1-C5), S7P(C1-C7)) as an input for our model (∑8 data sets) and disregarded Q4, P_Input_, S_Input_ and T_Input_ for model structure alignment. The following fluxes were considered in the validation: ΔGPI, ΔQ2, ΔTPI, Q11, Z3, ΔTKT1, ΔTKT2, and ΔTAL (∑8 fluxes).

#### 2.8.7 Evaluation of flux precision for monitored fragments

Crown *et al*. has suggested the implementation of a precision score (*P*) for comparison of flux precision (Crown et al, 2016). This score *p*_*i*_ can be calculated for individual fluxes:

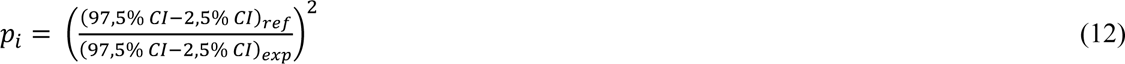

with CI referring to the confidence interval of a flux from an MFA only utilizing specific fragments as measurement input. These fragments can either be a reference fragment (ref, numerator) or fragments with precision scores of interests (exp, denominator). The precision score metric (*P*) was calculated as the average of individual flux precision scores (*p*_*i*_) for *n* selected fluxes:

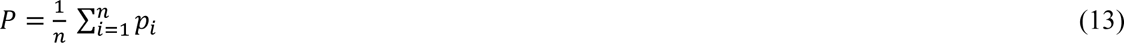

We selected the following fluxes of interest: Z3, ΔTAL, ΔTKT1, ΔTKT2, Q4, Q11, ΔGPI, ΔTPI, ΔQ2, ΔP_ex_, S7P input, and H6P input (endogen) (*n* = 12). A precision score greater than one was associated with increased flux precision compared to the reference experiment.

### 2.9 Principal component analysis

The PCA was performed with the ^13^C-MFA results obtained from 28 samples (resting granulocytes: n = 10, *E.coli*-stimulated granulocytes: n = 10, phorbol-12-myristate-13-acetate (PMA)-stimulated neutrophils: n = 5, PMA+diphenyleneiodonium chloride (DPI)-stimulated neutrophils: n = 3). The following estimated fluxes of interest were included in this analysis: Z3, Q4, S7P input, Q11, ΔQ2, ΔTAL, ΔTKT1, ΔTKT2, and ΔGPI. The ^13^C-MFA input data were standardized to a sample mean of zero and a unit sample standard deviation of 1 before performing the PCA in RStudio (version 2022.12.0) with the following built in R functions: prcomp, varimax (package stats), pracma, and fviz_eig (package factoextra) (Forina et al, 1989; R Core Team, 2020). We used the jackknife test, also called “leave one out” procedure for cross-validation (Abdi & Williams, 2010; Babamoradi et al, 2013; Efron & Tibshirani, 1986). Parameters (fluxes) were considered significant when the relative error (SE over nominal value) was smaller than 0.5. Finally, to interpret the fluxes, the PCA loadings were re-scaled to the original model values.

## 3 Results and discussion

### 3.1 GC-MS analysis of intracellular metabolites as their ethyloxime-trimethylsilyl derivatives

We used an ethyloxime-trimethylsilyl derivatization for determination of the ^13^C-labeling pattern of intracellular metabolites. This approach allowed us to obtain multiple GC-MS fragments containing labeling information of specific parts of the molecule along with the entire carbon skeleton (Table 1, Figure 2). Moreover, we achieved baseline separation of selected metabolites like glucose, DHAP, 3PG, and hexose-phosphate isomers (G1P, F6P, M6P, G6P) within a total run time of 15 min. Pentose phosphates (P5P) such as X5P, R5P, and Ru5P could not be clearly chromatographically separated, but electron ionization allowed the capture of specific labeling information of the lower three carbon atoms (C3-C5) of R5P. The acquired MID of the sugar phosphate fragments corresponded with the theoretical values, with an absolute error between 0.1 and 1 mol% (Supplementary file: research_data.xlsx.). Consequently, the monitored fragments were deemed suitable for ^13^C-MFA.

**Figure 2:**
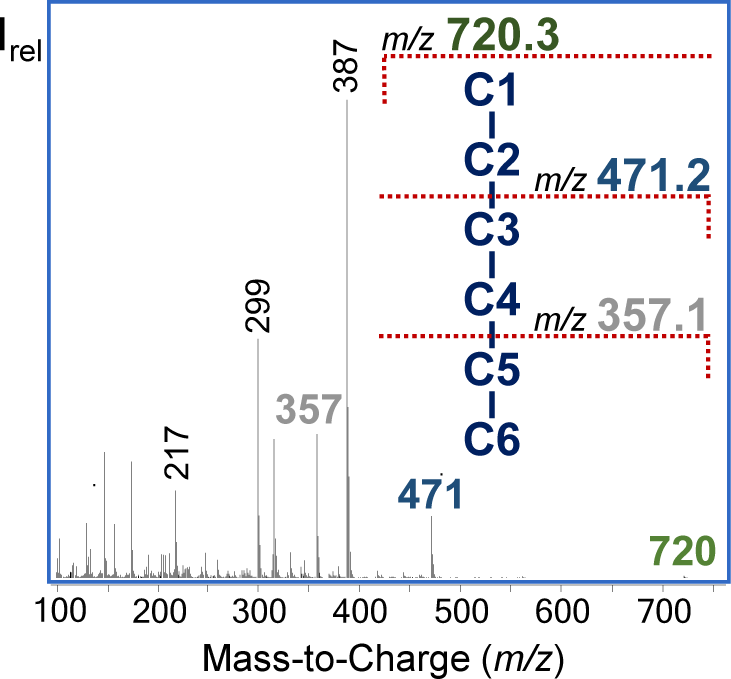
GC-EI-MS spectrum of G6P as an EtOx-TMS derivative (70eV).

**Table 1.** Selected GC-MS fragments as EtOx-TMS derivatives for ^13^C-MFA.

| Metabolite | $m/z$ | Carbon atoms | Molecular formula |
| --- | --- | --- | --- |
| G6P | 720 | C1 - C6 | $\text{C}_{25}\text{H}_{63}\text{O}_9\text{NSi}_6\text{P}$ |
| | 471 | C3 - C6 | $\text{C}_{16}\text{H}_{40}\text{O}_6\text{Si}_4\text{P}$ |
| | 357 | C5 - C6 | $\text{C}_{11}\text{H}_{30}\text{O}_5\text{Si}_3\text{P}$ |
| X5P, Ru5P, R5P | 618 | C1 - C5 | $\text{C}_{21}\text{H}_{53}\text{O}_8\text{NSi}_5\text{P}$ |
| R5P | 459 | C3 - C5 | $\text{C}_{15}\text{H}_{40}\text{O}_6\text{Si}_4\text{P}$ |
| DHAP | 414 | C1 - C3 | $\text{C}_{13}\text{H}_{33}\text{O}_6\text{NSi}_3\text{P}$ |
| 3PG | 459 | C1 - C3 | $\text{C}_{14}\text{H}_{36}\text{O}_7\text{Si}_4\text{P}$ |
| Glucose | 319 | C3 - C6 | $\text{C}_{13}\text{H}_{31}\text{O}_3\text{Si}_3$ |
| | 568 | C1 - C6 | $\text{C}_{22}\text{H}_{54}\text{O}_6\text{NSi}_5$ |

### 3.2 Evaluation of ^13^C-isotopomer mass distribution based on established models and concepts

#### 3.2.1 Characterization of glucose pathway utilization based on interpretation of ^13^C labeling patterns

Using [1,2-^13^C]glucose tracer experiments in combination with estimation of the M+1/M+2 enrichment ratio in downstream metabolites (e.g., 3PG) is the most widespread ^13^C labeling strategy to assess relative PPP activity. Metabolism of [1,2-^13^C]glucose through glycolysis generates M+0 and M+2 labeled trioses, while metabolism *via* oxidative PPP produces a combination of M+0, M+1 and M+2 labeled intermediates. Compared to untreated granulocytes we observed that the M+1/M+2 ratio in G6P, and 3PG increased more than twofold after stimulation with *E.coli* bioparticles (Figure 3a,c), indicating that phagocytosis leads to up-regulation of the oxidative PPP relative to the glycolytic pathway. However, Antoniewicz pointed out that the equilibration of G6P and F6P affects the M+1 and M+2 ratio, limiting this strategy for relative PPP utilization to a rough estimate (Antoniewicz, 2018). In this context, the notable M+1 fraction in G6P (C1-C6) provided strong evidence that PPP-derived [1-^13^C]F6P was channeled to G6P *via* glucose-6-phophate isomerase (GPI) and subsequently recycled into the PPP (Figure 3a). In consequence, this increased the M+0 and decreased the M+1 content, leading to an underestimation of PPP utilization. Moreover, the significant M+2 fraction in P5P (C1-C5) obtained after [1,2-^13^C]glucose tracer experiments indicated that P5P was produced *via* the non-oxidative PPP (Figure 3b). When observing the C3-C6 fragment of G6P, [1,2-^13^C]glucose-induced M+2 and M+1 fractions must have been transferred from C1 and C2 to the lower half of hexose-6-phosphate (H6P) (Figure 3f). Additionally, the high M+1 and M+3 fractions of the G6P fragment (C3-C6, *m/z* 471) after [U-^13^C] tracer experiments indicated that carbons from trioses were recycled to G6P (Figure 3d). Sadiku *et al*. detected multiple G6P isotopologues with a high M+3 fraction in neutrophils using [U-^13^C]glucose LC-MS tracing experiments, which were explained by the occurrence of gluconeogenesis to generate G6P (Sadiku et al, 2021). However, the M+3 labeling of G6P resulting from [U-^13^C]glucose tracer experiments can be induced by both the gluconeogenic condensation reaction of trioses as well as by the non-oxidative PPP *via* transaldolase-induced exchanges of GAP from the glycolytic pathway with carbons C4-C6 of F6P. Therefore, interpretation of this labeling data was challenging, as PPP and gluconeogenesis pathways share several metabolites like GAP, F6P, and G6P. We performed [4,5,6-^13^C]glucose tracer experiments for a rough assessment of the gluconeogenesis pathway, since, in contrast to the gluconeogenic condensation reaction, ^13^C labeled triose atoms would not be incorporated into the upper half of G6P through the non-oxidative PPP. When compared to the [U-^13^C]glucose tracer analysis we did not detect multiple isotopomers for the G6P fragment (*m/z* 471) (Figure 3d,e). In conclusion, we can exclude the presence of gluconeogenesis based on the absence of a M+1 and M+4, indicating no transfer of lower half to upper half carbons.

**Figure 3.**
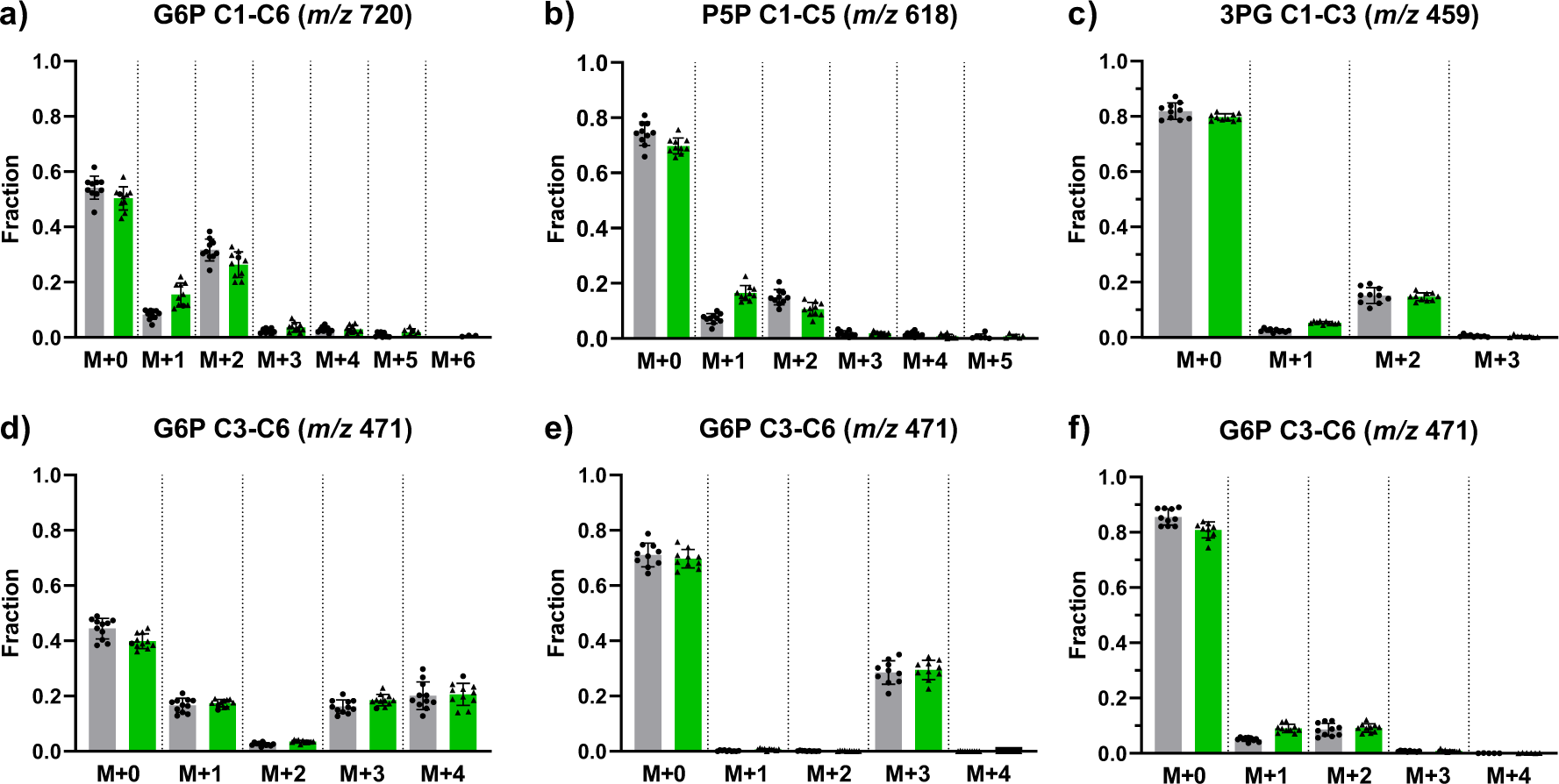
^13^C labeling patterns obtained from parallel tracer experiments with [1,2-^13^C]glucose (a-c, f), [U-^13^C]glucose (d) and [4,5,6-^13^C]glucose (e) of resting granulocytes (gray bars, black circle) and granulocytes after stimulation with E.coli bioparticles (green bars, black triangles). Mass isotopomer distributions were corrected for natural isotope abundance. Bar graphs are displayed as mean ± sd of n= 10 biological replicates.

#### 3.2.2 Incorporation of unlabeled reaction products

##### 3.2.2.1 Estimation of fractional contribution using the [U-^13^C]glucose tracer

The fractional contribution (FC) value resulting from fully labeled nutrients provides information about carbon sources contributing to the metabolite. In this respect, a decreased FC value may indicate incorporation of unlabeled reaction products (Buescher et al, 2015). Based on the mass isotopomer distribution, we calculated the FC as described above (2.8.2). The FC was significantly lower for G6P (*m/z* 720, C1-C6) compared to glucose (*m/z* 568, C1-C6), with a greater difference in untreated granulocytes compared to activated granulocytes (Figure 4a). Both glucose uptake *via* glucose phosphorylation by hexokinase and dilution through the release of unlabeled carbon sources like glycogen represent potential underlying causes for the overall lower FC of G6P. In this context, the [U-^13^C]glucose tracer analysis from Ma *et al*. demonstrated that LPS-stimulated macrophages utilized glycogen to generate G6P as a substrate for the PPP (Ma et al, 2020). Sadiku *et al*. further showed that the glycogen synthesis/degradation cycle is essential for neutrophil functions like phagocytosis (Sadiku et al, 2021). Both authors reported increased tracer enrichment in glycogen and hexose-6-phosphates combined with increased expression of glucose transporters upon activation of immune cells. In agreement with these findings, we observed a higher ^13^C enrichment of G6P upon activation of granulocytes, whereas the FC of glucose remained unaffected. Our data suggested that granulocytes increased hexokinase activity for bacterial killing, which is consistent with the finding of Ma *et al*. observing higher enzyme expression of hexokinase in IFN-γ/LPS-stimulated macrophages compared to untreated cells (Ma et al, 2020).

**Figure 4.**
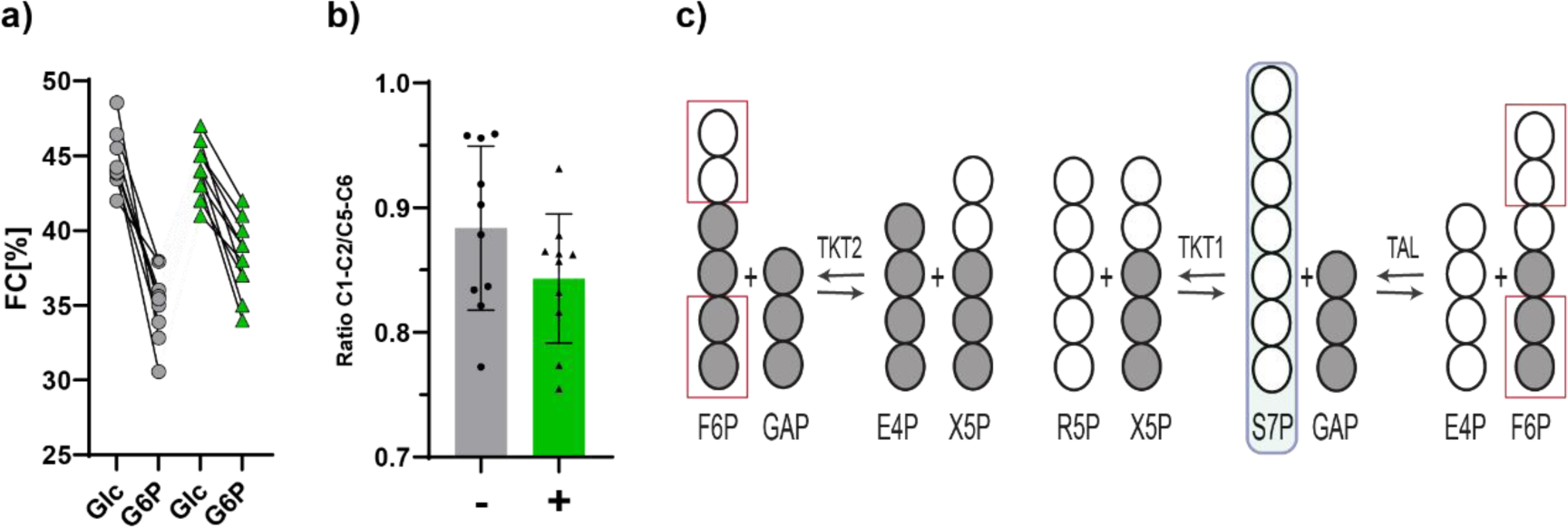
^13^C enrichment analysis of [U-^13^C]glucose tracer experiments. a) Fractional contribution (FC) of Glucose (Glc) (m/z 568) and G6P (m/z 720) from untreated (grey circle) and E.coli bioparticle-stimulated granulocytes (green triangle), n = 10. b) Ratio of total ^13^C enrichment in C1-C2 to the total ^13^C enrichment in C5-C6 of G6P obtained from untreated (gray bars, black circle) and E.coli-stimulated granulocytes (green bars, black triangle). Bar graphs are displayed as mean ± sd of n = 10 biological replicates. The individual values are indicated as dots. c) Reactions within the non-oxidative PPP: an unlabeled S7P input leads to increased ^13^C enrichment in the lower position (C5-C6) of H6P in contrast to the upper carbon atoms (C1-C2). Dark circles indicate ^13^C atoms, while the open circles represent ^12^C atoms.

##### 3.2.2.2 ^13^C-positional labeling analysis of G6P indicates an input of unlabeled S7P into the non-oxidative PPP

The results of the ^13^C-MID analysis indicated that both glycolysis as well as the non-oxidative PPP and the glycogen synthesis/degradation cycle contribute to the synthesis of G6P in granulocytes. Separate labeling data in the upper and lower half of G6P generally provides information to identify potential unlabeled carbon sources. In contrast to the ^13^C dilution of G6P through glycogenolysis and glycolysis, which affects the FC of the whole molecule, incorporation of unlabeled carbon sources in the PPP causes a decrease of ^13^C labeling in the upper carbon atoms of G6P (Figure 4c).

We aimed to establish a GC-MS method that provides ^13^C-postional labeling information of the upper G6P carbon atoms to address this effect. However, the method did not provide a suitable fragment containing only upper half carbons (C1-C2): although we identified a fragment *m/z* 176 containing C1-C2, it had to be dismissed for ^13^C-MID analysis due to peak overlap with another fragment of G6P in the same *m/z* range. As a workaround, we calculated the ^13^C enrichment of fragment C1-C2 by subtracting the total labeling of *m/z* 471 (C3-C6) from *m/z* 720 (C1-C6) (Equation 5). Both untreated and stimulated granulocytes displayed a lower ^13^C enrichment in the C1-C2 unit in contrast to the C5-C6 unit (*m/z* 356) with no significant intergroup differences (Figure 4b). These findings suggested an incorporation of unlabeled metabolites like S7P into the PPP. Interestingly, Haschemi *et al*. identified a seduheptulose kinase (CARKL) in macrophages that directs the metabolic state of the non-oxidative PPP by generation of S7P from seduheptulose (Haschemi et al, 2012). Another enzyme, sedoheptulose-1,7-bisphosphatase (ShB17), promoted R5P production in yeast without affecting oxidative PPP reactions by facilitating the generation of S7P from sedoheptulose-1,7-bisphosphate (Clasquin et al, 2011). Through similar mediators, the transfer of the upper two carbons of unlabeled S7P to GAP could potentially result in a reduced FC value of hexose-6-phosphates, especially in C1-C2 (Figure 4c).

### 3.3 Validation and evaluation of our Bayesian ^13^C-MFA performance

The ^13^C labeling data of sugar phosphates resulting from parallel tracer experiments of granulocytes were applied to ^13^C-MFA. The proposed reaction network is introduced in Figure 1. Its structure was built on the basis of the Katz/Rognstad model (Katz & Rognstad, 1967). For comparability with the recent literature report from Britt *et al.,* we included separate F6P and G6P compartments and an unlabeled glucose input. We also estimated 6PG and F-1,6-B patterns from the patterns of other metabolites in our system (see Bayes_implementation.pdf). In addition, the above-mentioned results (3.2.2.2) of our ^13^C-positional labeling analysis of G6P (Figure 4) indicate an input of endogenous carbon sources into the PPP. Therefore, we considered an input of unlabeled material into the P5P (P_Input_), S7P (S_Input_), and DHAP (T_Input_) pool. However, as the T_Input_ had no significant impact on the results, we disregarded it in our MFA routine.

#### 3.3.1 Validation of the proposed ^13^C-MFA approach: accuracy and precision were comparable with state-of-the-art software packages

In contrast to a Bayesian approach, the most common software packages for ^13^C-MFA use non-linear regressions or optimization approaches based on frequentists statistics to determine the fluxes, where confidence intervals are commonly either calculated with Monte Carlo analysis or parameter continuation (Antoniewicz et al, 2006; Young, 2014). For validation of our novel Bayesian ^13^C-MFA approach, we compared our results against a widely used INCA-software that was applied by Britt *et al*. (Britt et al, 2022). For this purpose, we used the same ^13^C-labeling data and model structure (modified as outlined in Section 2.8.4.3) and achieved comparable estimated flux values and precisions (Figure 5). The marginal differences between our and Britt *et al*.’s results can be traced back to a different handling of the measurement error and different upper and lower bounds for the fluxes. A list of flux results with their uncertainties of each donor are available in the Supplementary file: research_data.xlsx.

**Figure 5.**
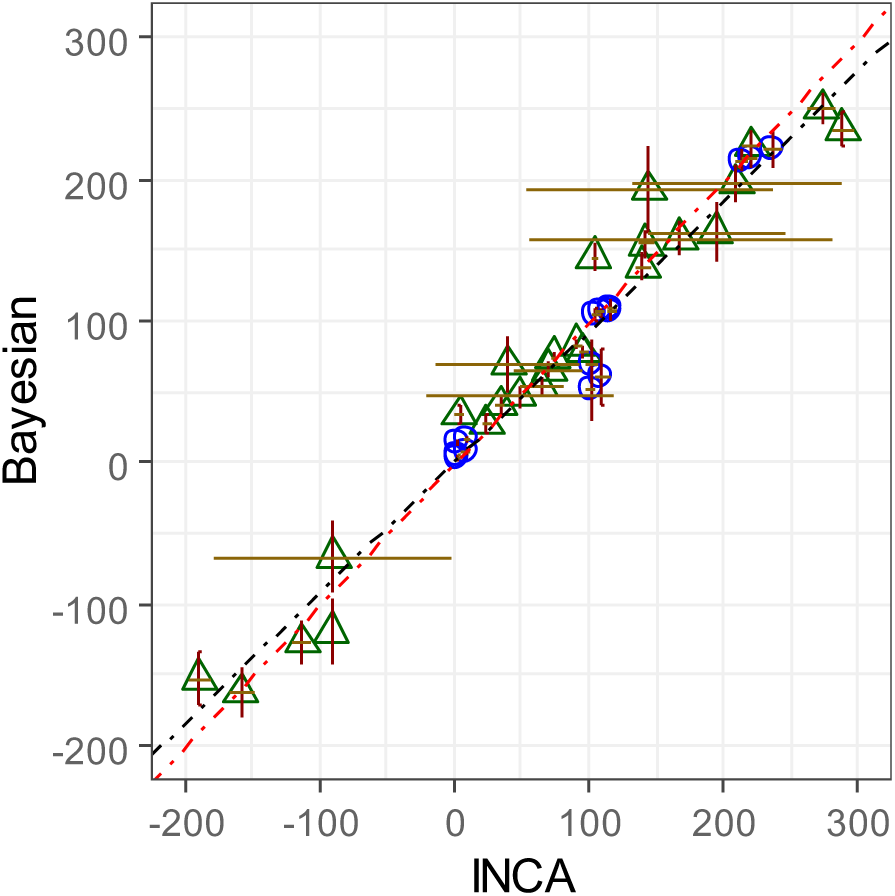
^13^C-MFA results of our Bayesian ^13^C-MFA in comparison with the ^13^C-MFA results published by Britt et al. estimated using INCA-software. Both MFA approaches utilized the same ^13^C labeling data. Symbols represent mean ± sd of individual fluxes (∑8) of 8 samples (∑64 data points). Green triangles: PMA-stimulated neutrophils, blue circles: PMA+DPI-stimulated neutrophils, brown error bars: horizontal, dark red error bars: vertical, red dotted line: y=x, black dotted line: linear regression y=0.92x+2.78 (R^2^ = 0.958).

#### 3.3.2 Selection of ^13^C-labeled glucose tracers for ^13^C-MFA of glycolysis and PPP

The selection of tracers for MFA is a critical step for accurate and precise flux determination (Antoniewicz, 2015). To investigate whether the selected tracer combination of [4,5,6-^13^C]glucose, [U-^13^C]glucose, and [1,2-^13^C]glucose in combination with our proposed GC-MS analysis improves flux resolution of our metabolic model, we tested different ^13^C-MFA scenarios. Firstly, we performed individual ^13^C-MFA for each tracer, followed by ^13^C-MFA with any combination of two tracers and a complete ^13^C-MFA by applying all data sets of parallel tracer experiments to the metabolic model. For the different ^13^C-MFA scenarios, synthetic data sets were utilized to exclude the effects of potential model and measurement errors from our error analysis. Simulated data sets were generated as a random sample of different Dirichlet distributions for calculated values of a real data set.

Each tracer had unique benefits: In accordance to current literature, the [1,2-^13^C]glucose tracer was the optimal single tracer setup to investigate the PPP (Figure 6), whereas the [U-^13^C]glucose tracer was the best performing tracer to determine tracer dilution. The [4,5,6-^13^C]glucose tracer experiment resulted in a narrow confidence interval for assessment of the condensation reaction of DHAP and GAP (QR) when compared to both [1,2-^13^C]glucose and [U-^13^C]glucose tracer. These findings confirmed our previous assessment that the [U -^13^C]glucose tracer is not suitable for quantification of gluconeogenic pathways. Overall, parallel tracer experiments with the selected tracer combination strengthened flux information and provided partially redundant measurements, leading to a significant improvement in flux resolution of the complex glucose metabolism by glycolysis and PPP.

**Figure 6.**
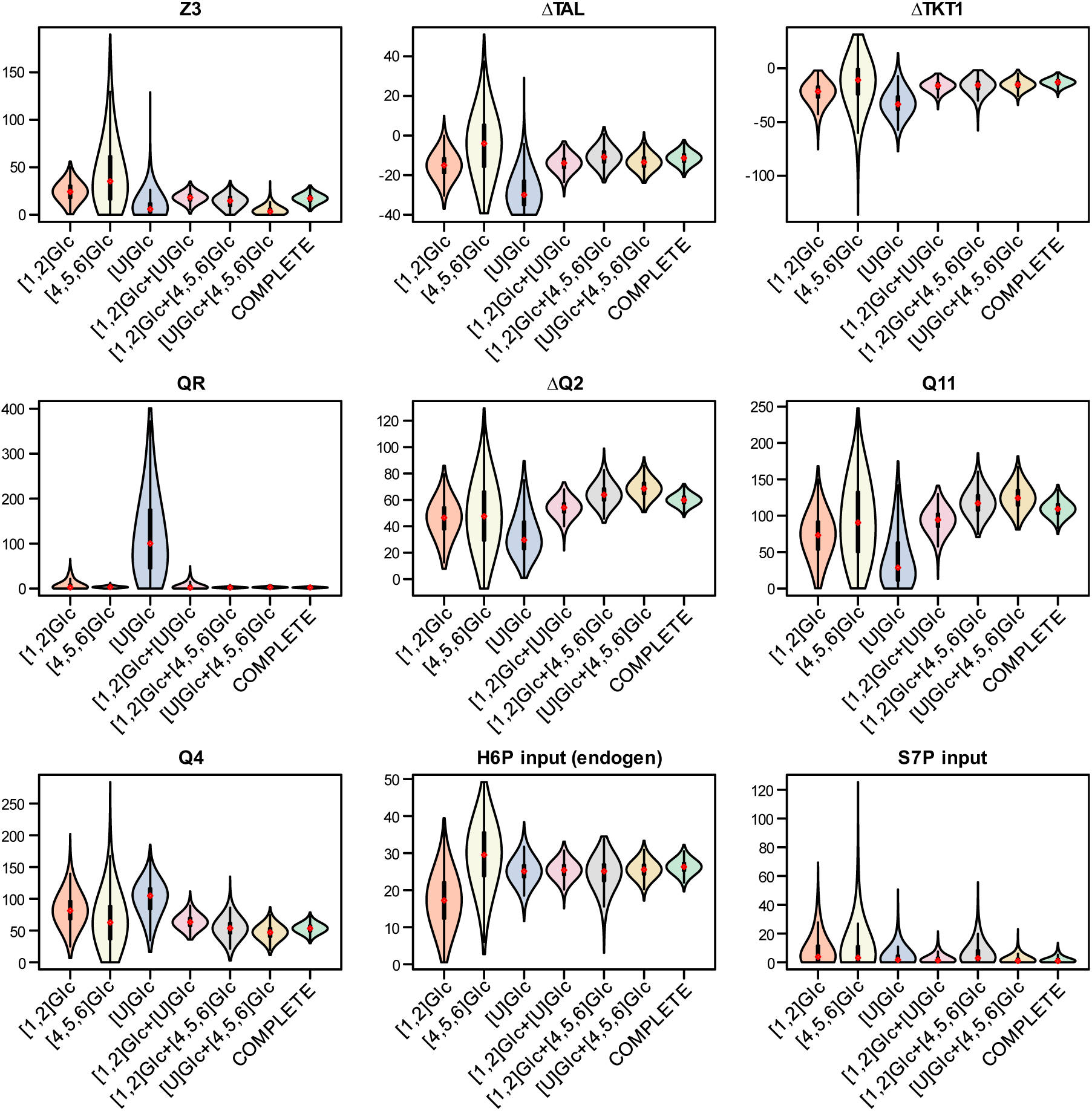
^13^C-MFA using all ^13^C labeling data (7 fragments, 4 metabolites) of individual tracer experiments and combined ^13^C-MFA of two to three tracers (complete ^13^C-MFA). Fluxes were normalized to a glucose uptake rate of 100. Results were obtained from a synthetic isotopomer data set. Violin plots represent a combination of box plot and kernel density plot. Median is highlighted as a white filled diamond with red border. Black bars represent the interquartile range (IQR) (first and third quartile). The lower/upper adjacent values are defined as first quartile -1.5 IQR or third quartile+ 1.5 IQR, respectively. They are visualized as thinner black lines extending from the IQR. The width of the violin plot reflects the frequency of data points. oxidative PPP: Z3; non-oxidative PPP: ΔTAL, ΔTKT1, Q4 (R5P loss); condensation of triose phosphates: QR; glycolytic fluxes: ΔQ2, Q11

#### 3.3.3 GC-MS fragments of G6P improve flux precision of the glucose metabolism

The quality of a ^13^C-MFA depends on information-rich ^13^C patterns. Labeling data of important sugar phosphates like G6P, one of the key metabolites of glucose metabolism, often remain elusive due to challenges in MS detection, while the application of labeling data from trioses like 3PG and DHAP is relatively widespread. Recently, improvement of PPP flux resolution was achieved by utilizing ^13^C labeling data obtained from GC-MS measurements of glucose and ribose after acid hydrolysis of glycogen and RNA (Long et al, 2016). However, our approach has proven less time consuming compared to this alternative approach, as sample preparation for our GC-MS analysis only required an extraction step of intracellular metabolites and the subsequent EtOx-TMS derivatization enabled the detection of several metabolites within a single GC−MS run. Compared to GC-NCI-MS and LC-MS techniques, our approach provided other fragments in addition to the fragment ion containing the intact carbon backbone of sugar phosphates, e.g., of G6P. Our final collection of fragments deemed suitable for ^13^C-MFA comprised 7 fragments of 4 intracellular metabolites. To estimate the advantage of using these fragments we performed the following ^13^C-MFA scenarios for the given parallel tracer experiments with [4,5,6-^13^C]glucose, [U-^13^C]glucose, and [1,2-^13^C]glucose: i) only 3PG labeling data as input, corresponding to the most commonly used PPP determination strategy, ii-iv): 3PG data in combination with either one of the three fragments of G6P, v) 3PG data combined with all three G6P fragments, vi) labeling data of 3PG in combination with DHAP and G6P (all fragments), and vii) all labeling data (∑7 fragments, 4 metabolites). To simulate these scenarios, we used synthetic data sets as previously described (3.3.2). Due to the EI fragmentation, the measurement precision of G6P fragment C1-C6 was lower when compared to fragment C5-C6 and C4-C6. To estimate the effect of the lower measurement precision, we included one additional scenario (G6P(C1-C6)*/3PG) comprising of the G6P (C1-C6) fragment but with the same precision values as defined for the other G6P fragments. Due to the greater precision of G6P (C1-C6), this scenario could provide measurements comparable to LC-MS or GC-NCI-MS.

The addition of G6P labeling information significantly increased the confidence in all flux results of glucose metabolism, with fragments of the lower half (C3-C6) being more informative compared to fragments C1-C6 and C5-C6 (Figure 7, Figure 8). Despite the low precision of G6P (C1-C6) measurements, its inclusion into the ^13^C-MFA analysis significantly enhanced flux precision, while the G6P fragment C5-C6 only lead to a marginally improved precision when compared to using only 3PG labeling data (precision score 1.89 and 1.12, respectively). Higher precision of G6P fragment C1-C6 resulted in even narrower confidence intervals. However, the performance with all three G6P fragments increased the precision score about two times compared to improvements through higher measurement precision due to simultaneous fitting of all three G6P fragments providing redundant information. Additional labeling information of DHAP combined with 3PG and G6P only slightly further improved both glycolytic and PPP flux determination. Drastic improvement in precision was observed for the condensation reaction of triose (QR). The labeling data of P5P (C1-C5, C3-C5) greatly improved flux resolution of the PPP but hardly provided any information about glycolytic fluxes like Q11 and ΔQ2.

**Figure 7.**
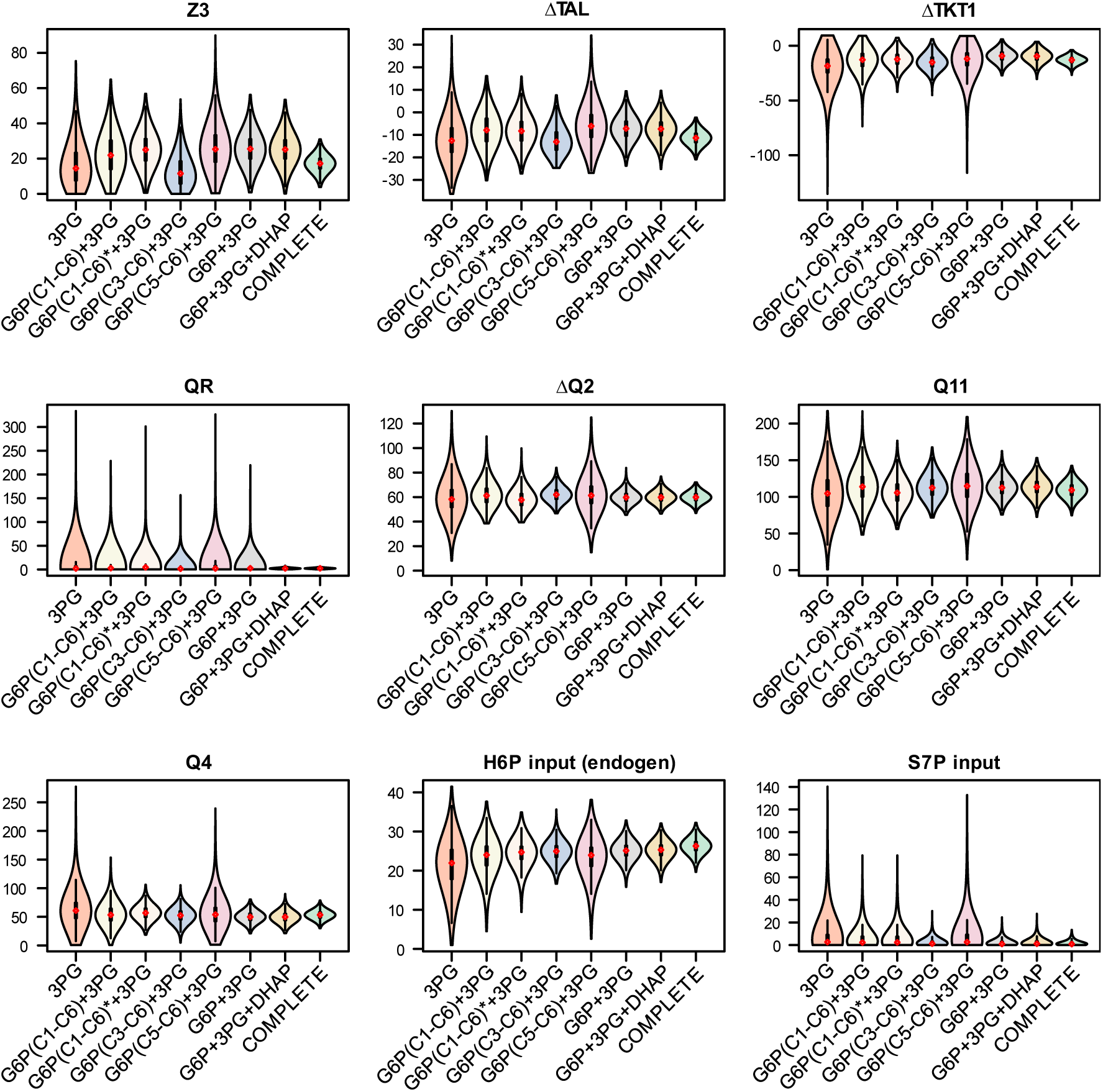
^13^C-MFA with 3PG labeling data (C1-C3) in comparison with i) additional G6P labeling data of individual fragments (C1-C6, C3-C6, C5-C6). The star symbol in the set description indicates that the measurement precision of G6P(C1-C6) was set to a value equivalent to the GC-MS precision of G6P fragments C3-C6 and C5-C6 ii) a combination of all three G6P fragments additionally to 3PG iii) a combination of 3PG, DHAP and G6P labeling data and iv) a complete MFA (3PG, DHAP, P5P, G6P ∑7 fragments). Results were obtained from a synthetic isotopomer data set derived from parallel tracer experiments with [4,5,6-^13^C]glucose, [U-^13^C]glucose, and [1,2-^13^C]glucose. Fluxes were normalized to a glucose uptake rate of 100. Violin plots represent a combination of box plots and kernel density plots. Median is highlighted as a white filled diamonds with red border. Black bars represent the interquartile range (IQR) (first and third quartile). The lower/upper adjacent values are defined as first quartile -1.5 IQR or third quartile+ 1.5 IQR, respectively. They are visualized as thinner black lines extending from the IQR. The width of the violin plot reflects the frequency of data points. oxidative PPP: Z3; non-oxidative PPP: ΔTAL, ΔTKT1, Q4 (R5P loss); condensation of triose phosphates: QR; glycolytic fluxes: ΔQ2, Q11

**Figure 8.**
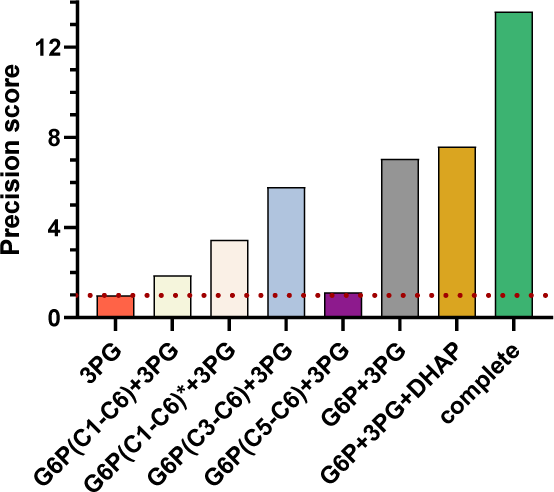
Precision scores for the given parallel tracer experiment ([1,2-^13^C]glucose, [U-^13^C]glucose, and [4,5,6-^13^C]glucose) while using different metabolites/fragments for ^13^C-MFA. The precision scores of the reference experiment (only 3PG labeling data) is 1 by definition and indicated as a red dotted line. The star symbol in the set description indicates that the measurement precision of G6P(C1-C6) was set to a value equivalent to the GC-MS precision of G6P fragments C3-C6 and C5-C6.

Overall, the confidence interval of the glucose metabolism, i.e. glycolysis and PPP was improved approx. 14-fold when labeling data of the selected GC-MS fragments (G6P (C1-C6, C4-C6, C5-C6), DHAP, 3PG, and P5P (C1-C5, C3-C5)) were included in ^13^C-MFA in contrast to the reference experiment (3PG only) (Figure 8).

### 3.4 Glucose metabolism of granulocytes: Identification of metabolic patterns and their regulation

#### 3.4.1 The non-oxidative PPP promotes ribose-5-phosphate biosynthesis in granulocytes

The ^13^C-MFA results showed that stimulation with *E.coli* bioparticles increased the oxidative PPP step (Z3) approximately 4-fold (Figure 9). In addition, the equilibrium between F6P and G6P was shifted towards G6P to enhance the oxidative PPP (Z3) to meet NADPH demand. This latter observation agrees at least partially with the reports by Britt *et al*. observing a reversed GPI net flux during oxidative burst of neutrophils (Britt et al, 2022). These findings supported our previously mentioned assumption that the Lee model leads to underestimation of the relative PPP activity (3.2.1). Moreover, the phagocytic process in granulocytes significantly shifted the transaldolase/transketolase reaction towards glucose catabolism and increased triose utilization (Q11). The constant ΔTPI flux suggests that PPP-derived GAP was channeled towards the glycolytic pathway. Interestingly, we found an S7P input under resting conditions, suggesting that uptake of unlabeled S7P into the non-oxidative PPP promotes R5P production for nucleotide biosynthesis (Q4). The latter combined with the S7P uptake decreased significantly upon stimulation, indicating a shift away from an R5P release. Our results therefore underscore the presence of a novel mechanism impacting glucose metabolism through the release of carbon sources into the non-oxidative PPP, adding further weight to the hypotheses proposed by Haschemi and others (Haschemi et al, 2012; Nagy & Haschemi, 2015; Ying et al, 2012).

**Figure 9.**
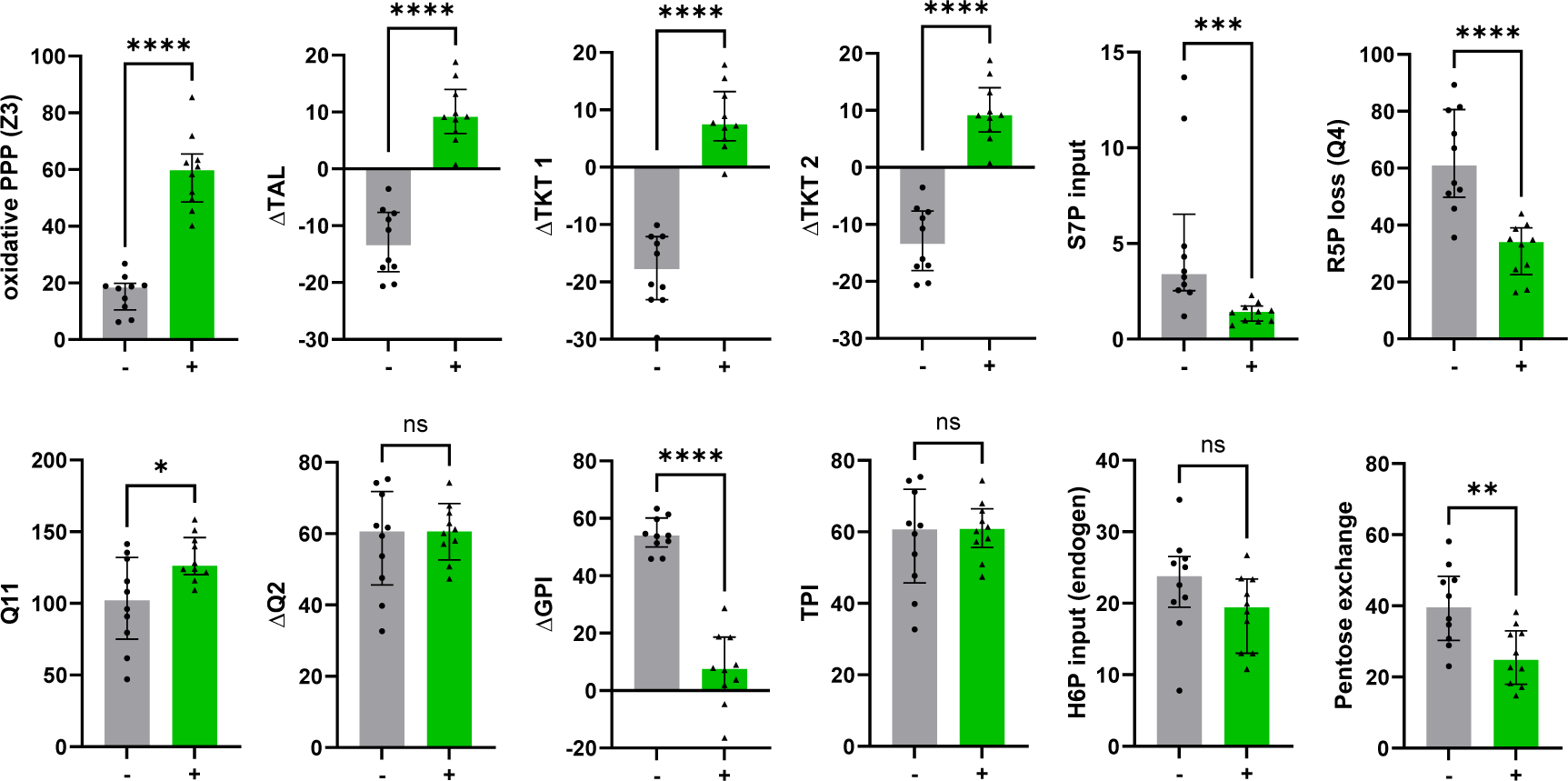
^13^C-MFA results of untreated granulocytes (-) (gray bars, black circles) and granulocytes stimulated with E.coli bioparticles (+) (green bars, black triangles). Fluxes were normalized to a glucose uptake rate of 100. Bar graphs show median with interquartile range of each group (n = 10). Individual symbols show posterior mean from Bayesian ^13^C-MFA. Statistical analysis was performed with the Mann-Whitney test (****= p < 0.0001, *** = p = 0.0003, ** = p = 0.0089, * = p = 0.0232).

#### 3.4.2 A “hard wired” PPP network gives insight into cellular mechanisms

The ^13^C-MFA results showed a strong inverse relation between ΔGPI and Z3 as well as a direct relation between ΔTAL and ΔTKT1 (Figure 10). This was the case for both i) data from MFA analysis of ^13^C labeling patterns that were obtained from our parallel tracer experiments and subsequent GC-MS measurements of granulocytes, and ii) data obtained from applying our ^13^C-MFA routine to LC-MS data of granulocytes from the recent publication of Britt *et al*.. Intriguingly, the relation between i) the net glycolytic fluxes ΔQ2 and Q11, ii) the R5P loss (Q4) and ΔQ2, and iii) ΔQ2 and ΔTAL were lost once neutrophils/granulocytes became stimulated. In contrast, we observed a relation between Z3 and ΔTAL upon immune cell activation. The frequently observed linear behavior between several fluxes could be partially explained by the stoichiometric restraint matrix (equation 11): 8 fluxes were calculated in respect of 4 - 6 independent fluxes (depending on which input fluxes were included), while Z1 was set to a fixed value of 100. Furthermore, the second and third column of the dependency matrix indicated that both ΔTAL and the glycolytic flux (ΔQ2) were closely linked to all other fluxes *via* constraints.

**Figure 10.**
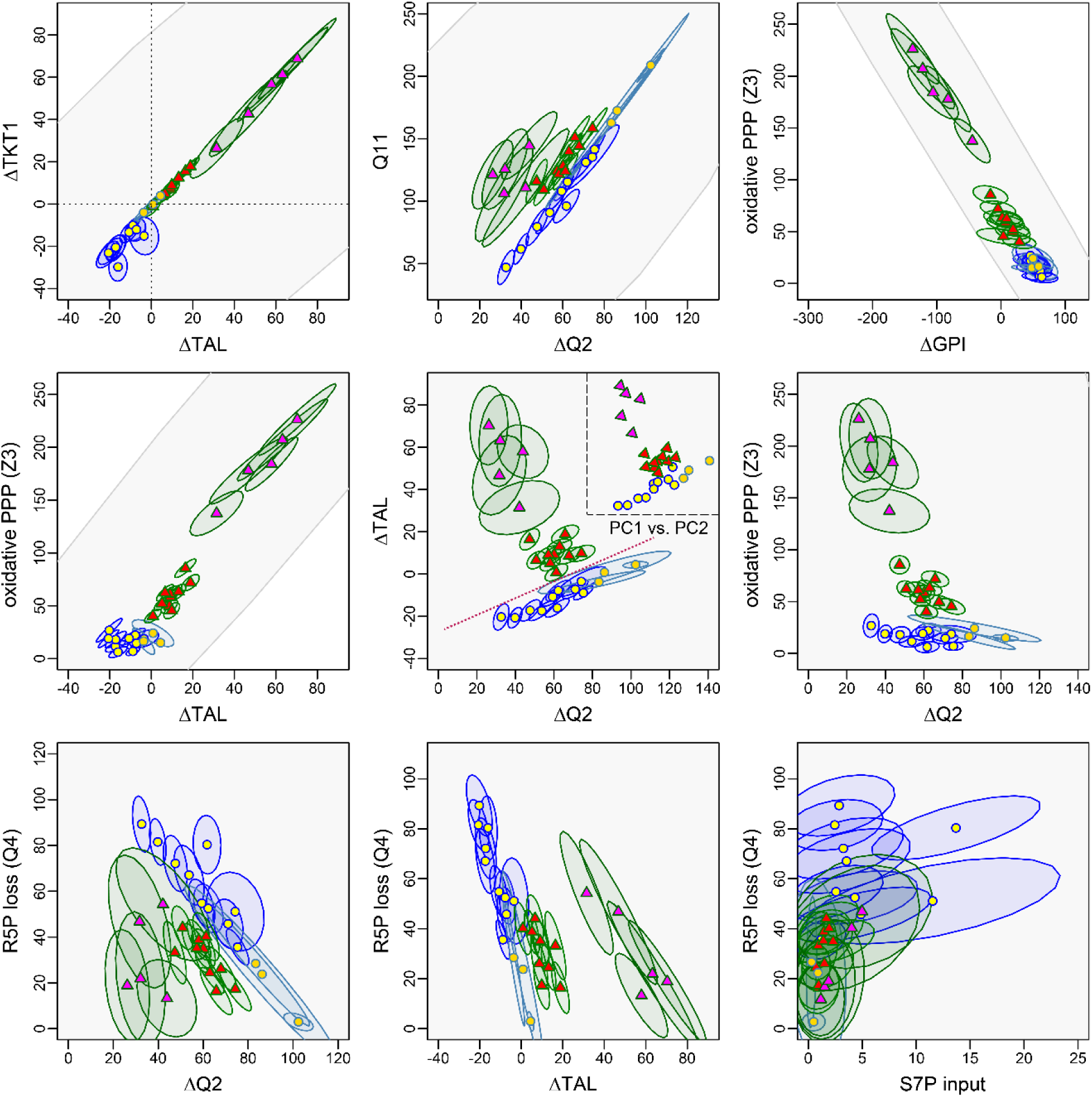
Relations between fluxes of the upper glucose metabolism. Results of the ^13^C-MFA are displayed as an ellipse with 0.68% confidence interval: dark blue ellipses with yellow circles indicate untreated granulocytes (n=10), green ellipses with red triangles granulocytes stimulated with E.coli bioparticles (n=10), light blue ellipses with beige circles PMA+DPI-stimulated neutrophils (n=3), and green ellipses with magenta triangles PMA-stimulated neutrophils (n=5). Symbols indicate the posterior mean of ^13^C-MFA datasets. ^13^C-MFA of granulocytes was performed with ^13^C labeling data obtained from parallel tracer experiments and subsequent GC-MS measurements. Neutrophil data were obtained by applying our ^13^C-MFA routine to LC−MS data from the recent publication of Britt et al.. The gray area represents the valid range based on the model structure without fitting of ^13^C labeling data. In the center graph at the top right, we added the PCA score plot of the first two principal components for comparison. The red dotted line indicates the separation between stimulated granulocytes and non-activated granulocytes.

Correlations between parameters can best be investigated with a PCA, which captures similar behaviors across several variables and thus allows to detect different directions of change. Interestingly, the PCA of the ^13^C-MFA results showed that three linear combinations of variables explain 99.6% of the variation in the data, with 86.7% of the variance being found in the first two principal component (PC1/PC2) axes. Figure 11 gives an overview over the first three PCs and their different regulatory processes (Figure 11; Table 2). PC1 referred to the directional shift in the PPP from quiescence to immune cell activation. In the positive direction, the flux from F6P to G6P increased together with the oxidative PPP. The resulting enhanced R5P production promoted the recycling of R5P to F6P and GAP instead of biosynthesis *via* the R5P loss of our system by either inhibiting the further use of R5P (e.g., for biosynthesis) or by an active shift in the reaction balance of transketolases. In addition, the GAP production from F6P was reduced. Analogously to results from Britt *et al*., the intense increase in oxidative PPP suggested a concomitant stimulation of this pathway by both enhanced NADPH demand and glycolytic backlog mediated by the GPI net flux (Britt et al, 2022). In summary, PC1 referred to the coupling of both PPP branches with the reversible glycolytic steps (ΔGPI, ΔQ2) to promote pentose cycling for maximization of the oxidative PPP or, in the negative direction, to the promotion of R5P synthesis by inhibition of the oxidative PPP. Both mechanisms did not affect pyruvate/lactate production. The second component indicated the linkage of non-oxidative PPP fluxes and downstream glycolytic fluxes to either maximize glycolytic pathways for energy supply or enhance R5P production for biosynthesis. This mechanism manifested in a positive net flux from F6P to R5P and a corresponding increase in R5P release. Furthermore, PC2 was completely independent of the oxidative PPP. This behavior could be explained by a regulatory shift of the non-oxidative PPP equilibrium towards R5P release. The associated consumption of glucose lead to a decrease in glycolytic triose and lactate production. The spread of data along the PCs 1 and 2 suggested a cell type and stimulation-specific degree of pentose phosphate production required for cell growth and function. In contrast, PC3 seemingly referred to the presence of cells in the sample that incorporated non-PPP-derived S7P into the non-oxidative arm of the PPP to promote pentose phosphate production.

**Figure 11.**
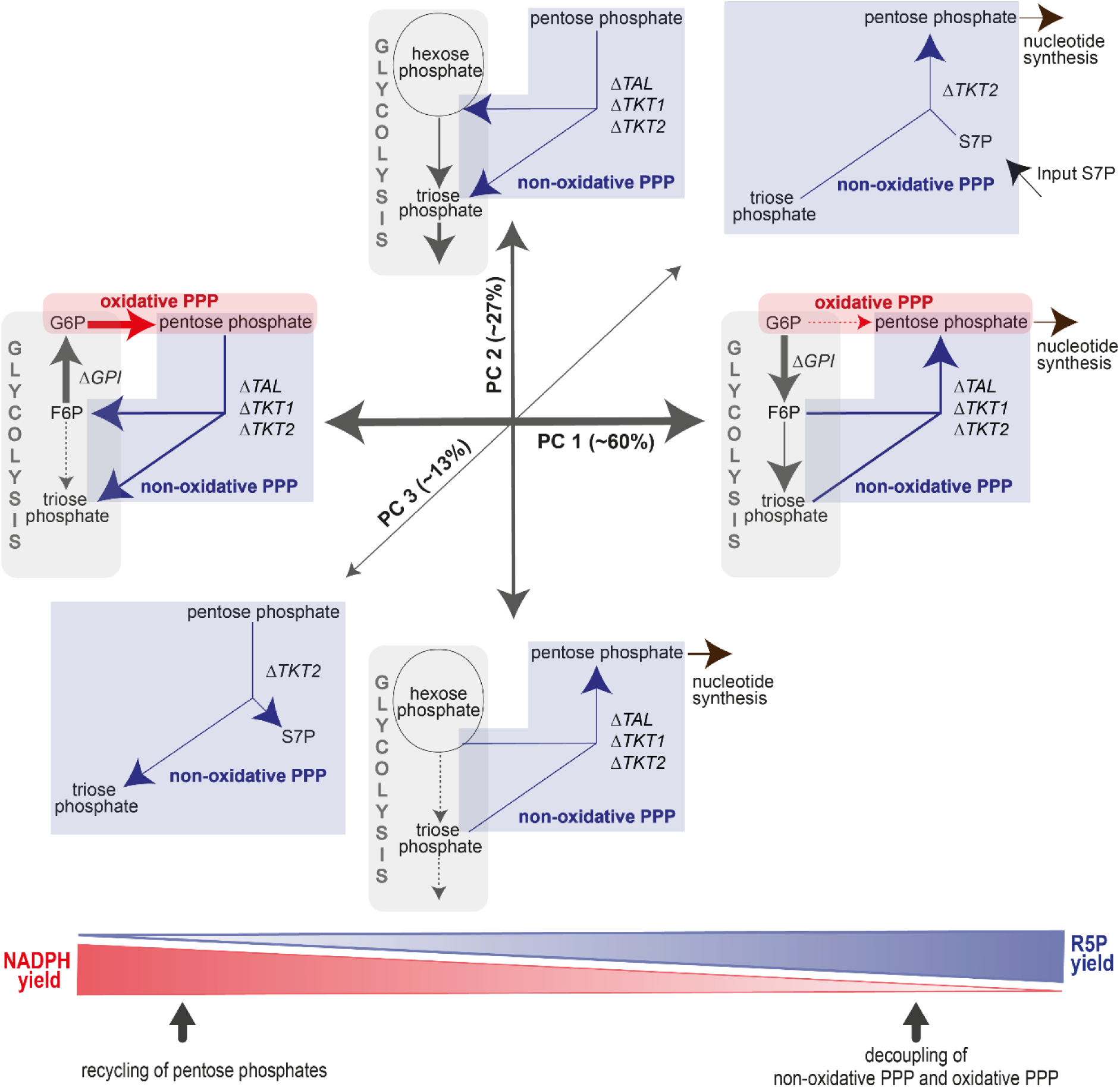
Metabolic states of granulocytes/ neutrophils identified by the first three principal components (PC) of PCA exploring ^13^C-MFA data (n=28, ∑_fluxes_ = 9). Contribution of each PC to the total variance indicated within parenthesis. The first three PCs amount to ∼99.6% of the variance.

**Table 2.** Metabolic fluxes that contributed significantly to the first three principal components (PC) of the PCA. Contribution of each PC to the total variance is indicated within parenthesis. The first three PCs amount to ∼99.6% of the variance. The sign indicates the direction of the PC relative to the mean.

| Metabolic flux |  | PC1 (60%) | PC2 (27%) | PC3 (13%) |
| --- | --- | --- | --- | --- |
| Glycolysis | ΔGPI | -57.2 | | -6.6 |
| | ΔQ2 | -10.0 | -15.0 | |
|  | Q11 |  | -32.7 |  |
| oxidative PPP | Z3 | 63.1 |  |  |
| non-oxidative PPP | ΔTAL | 24.4 | -3.2 | |
| | ΔTKT1 | 24.8 | -3.8 | 5.2 |
| | ΔTKT2 | 24.4 | -3.2 | |
|  | Input S7P |  |  | -3.0 |
|  | R5P loss (Q4) | -10.1 | 17.7 | -8.0 |

When plotting the main components PC1 and PC2 against each other, it is intriguing that they mirrored the scatterplot between ΔTAL and ΔQ2, suggesting that these key fluxes may regulate the glucose metabolism of granulocytes (Figure 10). However, it remained an open question whether these changes in ΔTAL were directly caused or simply passive responses to other regulatory processes. The stimulated and non-stimulated cells could be clearly separated with a line when assigning PMA+DPI-treated neutrophils to the unstimulated group (Figure 10), as the confidence regions for the flux determinations of the individual cells did not protrude into the other assignment region. The status of a single cell could thus be determined unambiguously.

## 4 Conclusion

In this study, we present a novel Bayesian ^13^C-MFA approach for determination of the PPP and glycolytic fluxes by utilizing ^13^C labeling data of various metabolic fragments including sugar phosphates in a parallel tracer experiment with [1,2-^13^C]glucose, [U-^13^C]glucose, and [4,5,6-^13^C]glucose. The 19 individual fluxes included in our glycolysis/PPP model could not independently assume all possible values as flux equilibrium had to be valid for each metabolite. A theoretical analysis *via* a stoichiometry restraint matrix showed that its permissible space could be reduced to three dimensions. This on one hand allowed an efficient and robust MCMC sampling for the Bayesian analysis and on the other hand revealed that the movement of all fluxes was limited to a three-dimensional subspace. Due to the reduced degrees of freedom, stimulation could be clearly separated from non-activated granulocytes. For this analysis, pairwise confidence ranges were necessary for the net transaldolase (ΔTAL) *vs* net glycolytic flux (ΔQ2) determinations which could only be determined *via* a Bayesian approach (with reasonable effort). Estimated fluxes showed strong correlation among themselves, with individual values moving on a single line in several cases. These frequently observed behaviors enabled a PCA analysis that detected three distinct axes of coordinated flux changes that were sufficient to explain all observations, in agreement with our theoretical analysis. Stimulation of granulocytes can be explained by the following mechanisms: Activation of oxidative PPP and inhibition of glycolytic F6P degradation to trioses occur in parallel. These steps are supported by inhibition of R5P release for biosynthesis, leading to enhanced carbon rearrangement of the non-oxidative PPP to form F6P for recycling into the oxidative PPP. Granulocytes in the resting state have enhanced R5P production, which can be explained by a regulatory shift of the non-oxidative equilibrium towards R5P. Metabolic changes of all analyzed cell experiments followed these behavioral/metabolic patterns irrespective of their cell population, stimulation intensity, and cell isolation method.

## Supporting information

EMU_Levels

Bayes_implementation

research_data

## Acknowledgements

This work was supported by the Deutsche Forschungsgemeinschaft (DFG, German Research Foundation, project 251293561 – collaborative research center CRC 1149), and the research training group GRK2203 PULMOSENS (Ulm University).

We are deeply grateful to Vittoria Passarelli, Rosemarie Meyer, Bettina Stahl, and Tanja Schulz for their excellent technical assistance.

## Supplementary material captions

### EMU_level.xlsx

The EMU approach calculates labeling on metabolites in successive levels, starting with labeling on isolated carbons and ending with the highest possible number of carbons on metabolites. Each level contains a linear system of equations with an inhomogeneous vector capturing input from outside and condensation products of labeled metabolite segments calculated in lower levels. This Excel file contains the matrices and input terms for levels 1 to 4.

### Bayes_implementation.pdf

This document introduces essential steps to implement the PPP/glycolysis model in the Stan Statistical Programming Language including i) definitions of appropriate priors such that sampling of parameters always yields non-negative fluxes. They are only subject to the constraint that a flux balance of zero results for each metabolite. ii) A simple model for the measurement error of ^13^C isotopomer distribution elements which provides error coupling between the individual distribution elements. iii) A reduction in the size of the network for efficient computation without introducing any bias in the calculated labeling.

### research_data.xlsx

This document provides the method parameters and datasets of our study: i) GC-MS settings including SIM parameters, ii) model parameters and their priors, iii) measured ^13^C-labeling patterns of granulocytes, iv) determined measurement errors of the ^13^C-labeling patterns, and v) flux results with the model-calculated labeling patterns and LogLik values.

## References

Abdi H, Williams LJ (2010) Principal component analysis. WIREs Comp Stat 2: 433–459

Antoniewicz MR (2013) 13C metabolic flux analysis: optimal design of isotopic labeling experiments. Curr Opin Biotechnol 24: 1116–1121

Antoniewicz MR (2015) Parallel labeling experiments for pathway elucidation and 13C metabolic flux analysis. Curr Opin Biotechnol 36

Antoniewicz MR (2018) A guide to 13C metabolic flux analysis for the cancer biologist. Exp Mol Med 50: 1–13

Antoniewicz MR, Kelleher JK, Stephanopoulos G (2006) Determination of confidence intervals of metabolic fluxes estimated from stable isotope measurements. Metab Eng 8: 324– 337

Antoniewicz MR, Kelleher JK, Stephanopoulos G (2007) Elementary metabolite units (EMU): A novel framework for modeling isotopic distributions. Metab Eng 9: 68–86

Babamoradi H, van den Berg F, Rinnan Å (2013) Bootstrap based confidence limits in principal component analysis — A case study. Chemometr Intell Lab Syst 120: 97–105

Backman TWH, Schenk C, Radivojevic T, Ando D, Singh J, Czajka JJ, Costello Z, Keasling JD, Tang Y, Akhmatskaya E et al (2023) BayFlux: A Bayesian method to quantify metabolic Flux es and their uncertainty at the genome scale. bioRxiv: 10.1101/2023.04.19.537435 [PREPRINT]

Borah Slater K, Beyß M, Xu Y, Barber J, Costa C, Newcombe J, Theorell A, Bailey MJ, Beste DJV, McFadden J et al (2023) One-shot 13C 15N-metabolic flux analysis for simultaneous quantification of carbon and nitrogen flux. Mol Syst Biol 19: e11099

Britt EC, Lika J, Giese MA, Schoen TJ, Seim GL, Huang Z, Lee PY, Huttenlocher A, Fan J (2022) Switching to the cyclic pentose phosphate pathway powers the oxidative burst in activated neutrophils. Nat Metab 4: 389–403

Buescher JM, Antoniewicz MR, Boros LG, Burgess SC, Brunengraber H, Clish CB, DeBerardinis RJ, Feron O, Frezza C, Ghesquiere B et al (2015) A roadmap for interpreting (13)C metabolite labeling patterns from cells. Curr Opin Biotechnol 34: 189–201

Carpenter B, Gelman A, Hoffman MD, Lee D, Goodrich B, Betancourt M, Brubaker MA, Guo J, Li P, Riddell A (2017) Stan: A Probabilistic Programming Language. J Stat Softw 76

Clasquin MF, Melamud E, Singer A, Gooding JR, Xu X, Dong A, Cui H, Campagna SR, Savchenko A, Yakunin AF et al (2011) Riboneogenesis in yeast. Cell 145: 969–980

Crown SB, Long CP, Antoniewicz MR (2015) Integrated 13C-metabolic flux analysis of 14 parallel labeling experiments in Escherichia coli. Metab Eng 28: 151–158

Crown SB, Long CP, Antoniewicz MR (2016) Optimal tracers for parallel labeling experiments and 13C metabolic flux analysis: A new precision and synergy scoring system. Metab Eng 38: 10–18

Efron B, Tibshirani R (1986) Bootstrap Methods for Standard Errors, Confidence Intervals, and Other Measures of Statistical Accuracy. Statist Sci 1

Forina M, Armanino C, Lanteri S, Leardi R (1989) Methods of varimax rotation in factor analysis with applications in clinical and food chemistry. J Chemom 3: 115–125

Hanke T, Nöh K, Noack S, Polen T, Bringer S, Sahm H, Wiechert W, Bott M (2013) Combined fluxomics and transcriptomics analysis of glucose catabolism via a partially cyclic pentose phosphate pathway in Gluconobacter oxydans 621H. Appl Environ Microbiol 79: 2336–2348

Haschemi A, Kosma P, Gille L, Evans CR, Burant CF, Starkl P, Knapp B, Haas R, Schmid JA, Jandl C et al (2012) The sedoheptulose kinase CARKL directs macrophage polarization through control of glucose metabolism. Cell Metab 15: 813–826

Hastings WK (1970) Monte Carlo sampling methods using Markov chains and their applications. Biometrika 57: 97–109

Jeon J-H, Hong C-W, Kim EY, Lee JM (2020) Current Understanding on the Metabolism of Neutrophils. Immune Netw 20: e46

Katz J, Rognstad R (1967) The labeling of pentose phosphate from glucose-14C and estimation of the rates of transaldolase, transketolase, the contribution of the pentose cycle, and ribose phosphate synthesis. Biochemistry 6: 2227–2247

Kuehne A, Emmert H, Soehle J, Winnefeld M, Fischer F, Wenck H, Gallinat S, Terstegen L, Lucius R, Hildebrand J et al (2015) Acute Activation of Oxidative Pentose Phosphate Pathway as First-Line Response to Oxidative Stress in Human Skin Cells. Mol Cell 59: 359–371

Kumar S, Dikshit M (2019) Metabolic Insight of Neutrophils in Health and Disease. Front Immunol 10: 2099

Li Y, Yao C-F, Xu F-J, Qu Y-Y, Li J-T, Lin Y, Cao Z-L, Lin P-C, Xu W, Zhao S-M et al (2019) APC/CCDH1 synchronizes ribose-5-phosphate levels and DNA synthesis to cell cycle progression. Nat Commun 10: 2502

Lima VF, Erban A, Daubermann AG, Freire FBS, Porto NP, Cândido-Sobrinho SA, Medeiros DB, Schwarzländer M, Fernie AR, Dos Anjos L et al (2021) Establishment of a GC-MS-based 13 C-positional isotopomer approach suitable for investigating metabolic fluxes in plant primary metabolism. Plant J 108: 1213–1233

Long CP, Au J, Gonzalez JE, Antoniewicz MR (2016) 13C metabolic flux analysis of microbial and mammalian systems is enhanced with GC-MS measurements of glycogen and RNA labeling. Metab Eng 38: 65–72

Ma J, Wei K, Liu J, Tang K, Zhang H, Zhu L, Chen J, Li F, Xu P, Liu J et al (2020) Glycogen metabolism regulates macrophage-mediated acute inflammatory responses. Nat Commun 11: 1769

Münz F, Wolfschmitt E-M, Zink F, Abele N, Hogg M, Hoffmann A, Gröger M, Calzia E, Waller C, Radermacher P et al (2023) Porcine blood cell and brain tissue energy metabolism: Effects of “early life stress”. Front Mol Biosci 10

Nagy C, Haschemi A (2013) Sedoheptulose kinase regulates cellular carbohydrate metabolism by sedoheptulose 7-phosphate supply. Biochem Soc Trans 41: 674–680

Nagy C, Haschemi A (2015) Time and Demand are Two Critical Dimensions of Immunometabolism: The Process of Macrophage Activation and the Pentose Phosphate Pathway. Front Immunol 6: 164

Okahashi N, Maeda K, Kawana S, Iida J, Shimizu H, Matsuda F (2019) Sugar phosphate analysis with baseline separation and soft ionization by gas chromatography-negative chemical ionization-mass spectrometry improves flux estimation of bidirectional reactions in cancer cells. Metab Eng 51: 43–49

Paclet M-H, Laurans S, Dupré-Crochet S (2022) Regulation of Neutrophil NADPH Oxidase, NOX2: A Crucial Effector in Neutrophil Phenotype and Function. Front Cell Dev Biol 10: 945749

Qi Liu, Fangming Zhu, Xinnan Liu, Ying Lu, Ke Yao, Na Tian, Lingfeng Tong, David A. Figge, Xiuwen Wang, Yichao Han et al (2022) Non-oxidative pentose phosphate pathway controls regulatory T cell function by integrating metabolism and epigenetics. Nat Metab 4: 559–574

R Core Team (2020) R: A Language and Environment for Statistical Computing, Vienna, Austria. Available from: https://www.R-project.org/

Rühl M, Rupp B, Nöh K, Wiechert W, Sauer U, Zamboni N (2012) Collisional fragmentation of central carbon metabolites in LC-MS/MS increases precision of 13C metabolic flux analysis. Biotechnol Bioeng 109: 763–771

Sadiku P, Willson JA, Ryan EM, Sammut D, Coelho P, Watts ER, Grecian R, Young JM, Bewley M, Arienti S et al (2021) Neutrophils Fuel Effective Immune Responses through Gluconeogenesis and Glycogenesis. Cell Metab 33: 411–423.e4

Simon-Molas H, Vallvé-Martínez X, Caldera-Quevedo I, Fontova P, Arnedo-Pac C, Vidal-Alabró A, Castaño E, Navarro-Sabaté À, Lloberas N, Bartrons R et al (2021) TP53-Induced Glycolysis and Apoptosis Regulator (TIGAR) Is Upregulated in Lymphocytes Stimulated with Concanavalin A. Int J Mol Sci 22: 7436

Stan Development Team (2020) RStan: the R interface to Stan. Available from: http://mc-stan.org/

Stanton RC (2012) Glucose-6-phosphate dehydrogenase, NADPH, and cell survival. IUBMB Life 64: 362–369

TeSlaa T, Ralser M, Fan J, Rabinowitz JD (2023) The pentose phosphate pathway in health and disease. Nat Metab 5: 1275–1289

Theorell A, Jadebeck JF, Nöh K, Stelling J (2022) PolyRound: polytope rounding for random sampling in metabolic networks. Bioinformatics 38: 566–567

Theorell A, Leweke S, Wiechert W, Nöh K (2017) To be certain about the uncertainty: Bayesian statistics for 13 C metabolic flux analysis. Biotechnol Bioeng 114: 2668–2684

Toller-Kawahisa JE, O’Neill LAJ (2022) How neutrophil metabolism affects bacterial killing. Open Biol 12: 220248

van Winden WA, Wittmann C, Heinzle E, Heijnen JJ (2002) Correcting mass isotopomer distributions for naturally occurring isotopes. Biotechnol Bioeng 80: 477–479

Wiechert W (2001) 13C metabolic flux analysis. Metab Eng 3: 195–206

Wolfschmitt E-M, Hogg M, Vogt JA, Zink F, Wachter U, Hezel F, Zhang X, Hoffmann A, Gröger M, Hartmann C et al (2023) The effect of sodium thiosulfate on immune cell metabolism during porcine hemorrhage and resuscitation. Front Immunol 14: 1125594

Ying H, Kimmelman AC, Lyssiotis CA, Hua S, Chu GC, Fletcher-Sananikone E, Locasale JW, Son J, Zhang H, Coloff JL et al (2012) Oncogenic Kras maintains pancreatic tumors through regulation of anabolic glucose metabolism. Cell 149: 656–670

Young JD (2014) INCA: a computational platform for isotopically non-stationary metabolic flux analysis. Bioinformatics 30: 1333–1335

Zamboni N, Fendt S-M, Rühl M, Sauer U (2009) (13)C-based metabolic flux analysis. Nat Protoc 4: 878–892

Zhang Y, Gao L (2003) On Numerical Solution of the Maximum Volume Ellipsoid Problem. SIAM J. Optim. 14: 53–76

