## Supplementary material for "Bayesian ^13^C-metabolic flux analysis of parallel tracer experiments in granulocytes: A directional shift within the non-oxidative pentose phosphate pathway supports phagocytosis": Bayes_implementation

### Adaptation of a $^{13}$C-MFA-Model to the *Stan* statistical language

**Abstract**

We adapted the t $^{13}$C-MFA approach to a Bayesian analysis by formulating the complete MFA-EMU [1,2] approach in the Statistical Programming Language 'Stan' [3–5]. In this supplement, we outline the implementation process of the MFA in Stan. In contrast to MFA implementations using nonlinear regression, Bayesian analysis is inevitably requires more calculation time but provides an exact distribution of the fluxes, their confidence regions, and their correlations. The cost of a numerical solution of equations grows with the third power of the number of metabolites included in the system. Therefore, there is an advantage to reducing the metabolites included in the reaction network to a 'core system'. Later, labeling patterns of non-included metabolites can be calculated from the core metabolites patterns without sacrificing model correctness.

The Bayesian approach uses a MCMC-sampling algorithm which draws random samples of a reservoir of unknown fluxes to assess the $^{13}$C labeling on various metabolites. Theses fluxes have to satisfy the restraints that the flux balances of all metabolite pools should be zero and no flux is negative. For flux sampling we define a permissible range that considers these restraints. The corresponding ranges allow initial distributions (priors) for the fluxes, that are non-informative and ensure that the model predictions are independent from the priors.

Self-developed programs may be erroneous. To minimize these errors, we introduced a number of 'self-consistency' tests.

#### Abbreviations

- *Hexoses*: F6P: fructose-6-phosphate; G6P: glucose-6-phosphate; 6PG: 6-phosphogluconate; F-1,6-B: fructose-1,6-biphosphate

- *Trioses*: GAP: glyceraldehyde-3-phosphate ; DHAP: dihydroxyacetone-phosphate
- *Pentoses*: Ru5P: ribulose-5-phosphate ; R5P: ribose-5-phosphate ; X5P: xylulose-5-phosphate
- *Sedupheptulose*: S7P: seduheptulose-7-phosphate
- *Erythrose:* E4P: erythrose-4-phosphate
- *CMD*: carbon mass distribution. We denote an element of this distribution as *isotopomer*, a superscript to a metabolite refers to the number of simultaneously labeled carbons, i.e. $G6P^3$.
- *EMU:* elementary metabolite unit

### 1 Bayesian parameter/flux estimation

For this multi-target optimization problem, we use a Bayesian analysis. With this analysis, the probability that a measured data set can be explained with a given model is optimized. To adapt this approach to our questions we denoted the set of all measured CMDs as the data $\mathbf{D}$ and the set of all unknown fluxes and other parameters is collected in the vector $\mathbf{q}$. Bayesian statistics defines this distribution as $\boldsymbol{P}(\mathbf{q}|D)$, the distribution of $\mathbf{q}$ given the data $D$, based on the relation [6]:

$$\boldsymbol{P}(\mathbf{q}|D) \propto \boldsymbol{P}(D|\mathbf{q})\boldsymbol{P}(\mathbf{q}). \tag{1}$$

In terms of the current application, $\boldsymbol{P}(D|\mathbf{q})$ represents the probability that the different predicted CMDs of each ion fragment for a given parameter/flux set $\mathbf{q}$ match the corresponding measured distributions. $\boldsymbol{P}(\mathbf{q})$ expresses the 'prior' knowledge about the parameter distributions.

The unknown parameters $\mathbf{q}$ are determined by optimizing the product of likelihoods defined in eqn (3), ideally leading to optimal $\mathbf{q}$ values. Slightly different parameters give a less optimal fit which in turn results in a distribution of parameter values whose shape depends on the prediction equation, measurement values and their error bounds The shape of their distribution $\boldsymbol{P}(\mathbf{q}|D)$ is generally unknown, though it can be estimated by using a sampling-based Markov chain Monte Carlo (MCMC) algorithm [7]. For implementation, we use the software package and statistical programming language 'Stan' [6]. Key elements of such an implementation comprise firstly the unknown parameters $\mathbf{q}$, secondly equations using these parameters to predict values of interest, and lastly 'sampling statements' that form the right side of eqn (1). In the underlying MCMC algorithm [7, 8], a parameter set is randomly selected and used to generate predictions that correspond to measured data. If the resulting predictions come close to the measurements in the frame of the measurement errors, then this sample is considered to be a valid sample of the underlying distribution and is collected in a sampling chain, otherwise it is disregarded. With this selection, the MCMC algorithm [7, 8] ensures that the distribution of the sampled parameters converges to the true underlying distribution with an increasing number of samples. Therefore, with a sampling run, one obtains a distribution for each parameter. The determination of an unknown parameter results in its distribution at the same time. The shape of this distribution depends solely on the model structure and the specification for the measurement error.

#### 1.1 Posterior values

Each time a new parameter set is added to the chain, a routine is started that uses the new, added parameter set to calculate other quantities, such as correction function values for fixed signal values. $^{13}C$ enrichment values from corrected MIDs are also possible. This creates a further sampling chain from which statistical properties of the additionally calculated quantities can be derived. This concept of 'posterior predictive sampling' is described in detail in the Stan user guide [9], chapter 28.1.

#### 1.2 The EMU approach

In this work, we applied the EMU approach [1] to $^{13}C$ mass distributions of metabolites. The EMU approach works with different levels, where each level considers labeling over neighboring carbons on metabolites of a reaction network. Level 1 works with abundance on isolated carbons, level 2 on mass distributions over two adjacent carbons, and level n refers to n-adjacent carbons. Labeling patterns of each level are calculated in order from level 1 to level n. For each level there is a linear system describing the relations of patterns and their loss from the network (input-output equations). An input can either consist of an input from outside of the network or of newly formed patterns calculated from the condensation of two different fragments, each of which originates from results of lower levels. With this strategy, mass distributions over complete molecules of a complex reaction network can be calculated via a series of linear equations without resorting to expensive iterative approximation methods. The data table `EMU_Level.xlsx` in the supplement contains the system of equations from level 1 to level 4 for our PPP model. It is noticeable that the size of the systems decreases with higher levels.

### 2 Measurement error handling

Metabolic fluxes are estimated by fitting model-calculated *CMDs* with corresponding distributions derived from mass spectrometry. This fitting should take the measurement error for the underlying signals into account. In the simplest case, it can be assumed that measurement errors for individual distribution elements are independent and proportional to the current value (about 5% of the nominal value). However, this simplification does not fully capture the current situation: when calculating a CMD, the signal area for a given isotopologue is divided by the sum of the areas of all isotopologues relevant to the distribution. The measurement errors of all signal areas thus have an influence on the denominator and induce a correlation between the individual distribution elements. Furthermore, it is not certain that relatively small distribution values have the same percent measurement error as large ones. Some metabolites are measured with greater signal strength than others and therefore with better precision. This dependence on the signal strength should be taken into account. To reflect this state, we use a Dirichlet distribution [10](Chp. 2.2.1). It describes the variation of a discrete distribution comprising of positive elements normalized to 1 that is often explained with the example of an urn scenario. In the urn there is a large number of k different balls with the distribution $[p_1, p_2, ..p_k]$. If n balls are drawn and afterwards put back into the urn, then we are given the following variance and covariance of elements of the distribution of collected balls:

$$\begin{aligned} \text{diagonal element: } var_i &= \frac{p_i(1-p_i)}{n+1} \\ \text{covariance element: } cov_{i,j} &= -\frac{p_i p_j}{n+1} \end{aligned} \tag{2}$$

As a result, one precision parameter is enough for a distribution. In the statistical modeling language, the likelihood that a measured distribution can be explained with the corresponding model calculation is:

$$meas_{G6P}[i] \sim dirichlet(pred_{G6P}[i] * prec_{G6P}); \qquad (3)$$

with $meas_{G6P}[i]$ as the mass distribution of the complete G6P fragment measured for the i-th tracer protocol, $pred_{G6P}[i]$ as the corresponding model prediction, and $prec_{G6P}$ as the appropriate precision. According to eqn (2) the standard deviation of a distribution element with value 0.1 and a Dirichlet precision of 1000 is 0.0095. The log-likelihood of a Dirichlet distribution captures the probability that the error in a measurement of a distribution can be explained by a given precision. For a molecular fragment with 6 carbons, the optimal log-likelihood would lie between 20 and 25. A still acceptable congruence between measured and theoretical distribution yields a log-likelihood value around 10. For a poor congruence it can become negative. In this case, the precision must be reduced so that the Dirichlet distribution can explain the large difference between the measured and theoretical distributions.

The natural m+1 enrichment for a 6 carbon fragment is about 0.06. If we used distributions corrected for natural $^{13}C$ labeling, the value would become essentially zero. In this case, the distribution error could not be defined. We therefore decided to use CMDs that are not corrected for natural $^{13}C$ abundance for distribution fitting, which might be less conservative within the metabolic community, but more useful for our precision estimation.

### 3 PPP-Modeling: The basic network

Figure 1 introduces the model used in the current project. The rudimentary structure was based on the Katz et al. model [11]. We explored how pentose moiety, generated by the oxidative part of the PPP, was either lost as R5P or channeled back to the hexose and triose pools, from where it may leave the system as pyruvate and lactate. The steps involved in this reaction sequence were the reversible flows of the non-oxidative PPP, namely

$$\begin{aligned} &\text{transaldolase} \quad \text{F6P+E4P} \underset{Q_9}{\overset{Q_{10}}{\rightleftharpoons}} \text{S7P+GAP} \\ &\text{transketolase I} \quad \text{X5P+E4P} \underset{Q_6}{\overset{Q_5}{\rightleftharpoons}} \text{F6P+GAP} \\ &\text{transketolase 2} \quad \text{X5P+R5P} \underset{Q_8}{\overset{Q_7}{\rightleftharpoons}} \text{S7P+GAP} \end{aligned} \tag{4}$$

The model was expanded to allow a negative net flow for these reaction pairs, such that i.e. $Q_{10}$ can be larger than $Q_9$. Moreover, by establishing $Q_R$ we considered a 'gluconeogenic' backflow from the triose pools to the hexose pool. The combined input into the G6P and F6P pools was set to 100, such that $F_{input}$ referred to the input into F6P and $100 - F_{input}$ to the G6P input. Both inputs can include unlabeled material. In addition, we allowed an input of unlabeled material into the pentose pool ($P_{Input}$), seduheptulose pool ($S_{input}$), and DHAP pool ($T_{input}$).

### 4 PPP model reduction to a core system

With the linearity of the equations for each EMU level, labeling patterns of a particular metabolite can be calculated from a superposition of the patterns on the supplying fluxes. Thus, some metabolites of the network, such as G6P or Ru5P, can be omitted, while metabolites representing central nodes are retained. The latter then form a core system. Given a solution of the core system, labeling patterns of an omitted metabolite can be calculated from a superposition of patterns of the core system.

Omitting some metabolites can significantly increase the efficiency in calculating the labeling patterns. For the PPP model shown in Fig.1, the number of isolated carbons in the system that can be labeled amounts to 34. In consequence, a 34 x 34 matrix must be inverted to calculate their enrichments. The number of elementary multiplication and addition steps required for the inversion of a matrix is proportional to the third power of the edge length of the matrix. In our case this would account to about 40000 elementary steps and therefore impose a large computational load, given the fact that the inversion step must be repeated thousands of times in the context of a Bayesian analysis. It is possible to deduct 11 carbons if the patterns of G6P and Ru5P are expressed by other patterns. Because of the dependence

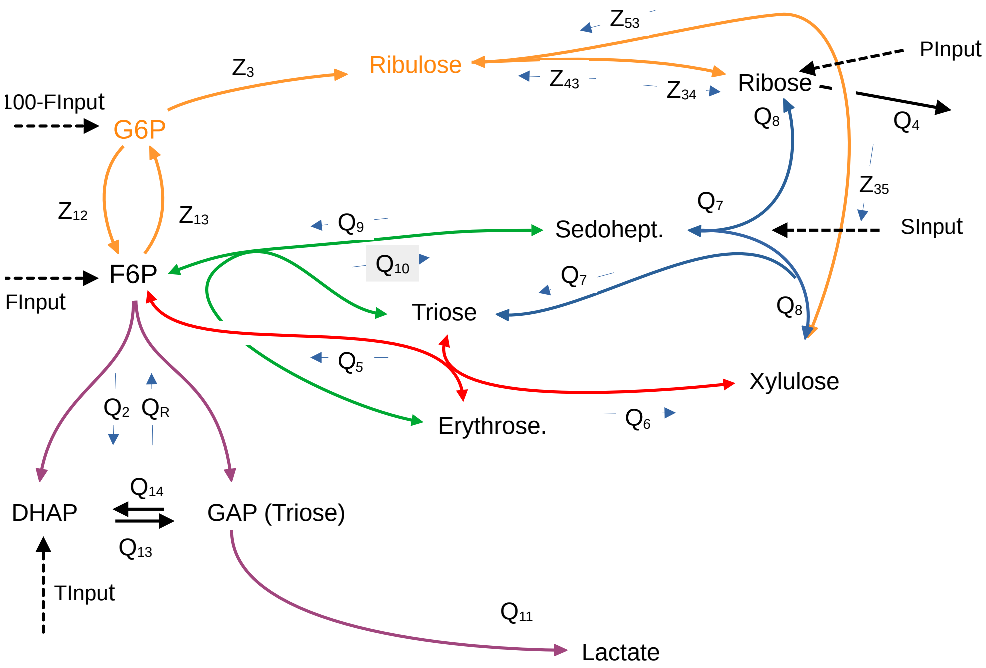


Figure 1: Flows of the glycolytic and PPP reaction network as implemented in the model. The color code for flows is as follows: oxidative PPP: orange; glycolysis: violet; non-oxidative PPP: transketolase reactions: blue and red; transaldolase reaction: green.

on the third power, the associated computational effort is reduced to about 12000 steps, which equals about one third of the total effort. This showcases why it is desirable to reduce the complete PPP system to a minimal PPP core that retains all essential information about flows and labeling patterns. Hence, we introduce details of this reduction of considered metabolites in the next section. In Fig. 1, flows which are to be replaced start with upper case Z, while the new, merged fluxes of the reduced system are denoted $Q_x$ like other fluxes of the core system.

To define the core system, the F6P and G6P pools are condensed to one hexose compartment. Furthermore, in the pathway from G6P to R5P and X5P the intermediate step through Ru5P will be omitted. The mass balances for the to be simplified subsystem are:

$$\begin{aligned}
&\text{Input} && = \text{Output} \\
G6P :& G_{input} + Z_{13} && = Z_{12} + Z_3 && \text{(5a)} \\
F6P :& F_{input} + Z_{12} + Q_5 + Q_9 + Q_R && = Z_{13} + Q_6 + Q_{10} + Q_2 && \text{(5b)} \\
\text{Ru5P} :& Z_3 + Z_{43} + Z_{53} && = Z_{35} + Z_{34} && \text{(5c)} \\
\text{R5P} :& Z_{34} + Q_8 && = Q_4 + Q_7 + Z_{43} && \text{(5d)} \\
\text{X5P} :& Z_{35} + Q_8 + Q_6 && = Q_5 + Q_7 + Z_{53} && \text{(5e)}
\end{aligned}$$

The left side of the equations lists all fluxes entering, and the right the ones leaving a compartment. Combined these fluxes represent the total throughput of the compartment. For the following we define:

$$\begin{aligned}
F_{Input} + G_{Input} =& Z_1 = 100 \\
Q_5 + Q_9 + Q_R = Q_{inp}; &\quad Q_6 + Q_{10} + Q_2 = Q_{loss} \\
flux_f = Z_{13} + Q_{loss}; &\quad flux_g = Z_{12} + Z_3 \\
flux_{Ru5} =& Z_{35} + Z_{34} \qquad (6)
\end{aligned}$$

Firstly, we consider the mass balances of carbon isotopomers with the mass offset m for 6-carbon metabolites of this subsystem. Their mole fractions are denoted as $T^m_{G6P}$ and $T^m_{F6P}$ for the tracer input, $Q^m_{inp}$ for the transfer of label on the combined input $Q_{inp}$ from the non-oxidative PPP, and $G^m$ and $F^m$ for the G6P and F6P metabolites. During the decarboxylation of G6P to Ru5P one labeling position is lost. To indicate that the exchange of isotopomers on a 5 carbon fragment between the Ru5P, R5P and X5P pools cannot be easily linked with isotopomers on a 6 carbon fragment, we use the mass offset p for the pentose subsystem. The transketolase reactions of $Q_6$ and $Q_8$ involve a condensation of two precursors. We use the notation $c(Q_8)^p$ to refer to the transfer of $Q_8[S_{1-2} \otimes GAP]^p$ with the mass offset p on the condensation product. Accordingly, $c(Q6)^p$ refers to $Q_6[H_{1-2} \otimes GAP]^p$. With this abbreviated notation, the mass balances for the isotopomers of the hexose/pentose subsystem are defined as follows:

$$\begin{aligned}
G_{input}\, T^m_{G6P} + Z_{13}\, F^m &= flux_g G^m && \text{(7a)} \\
F_{input} T^m_{F6P} + Z_{12}\, G^m + Q^m_{inp} &= flux_f F^m && \text{(7b)} \\
Z_3\, T^p_5 + Z_{43}\, \text{R5P}^p + Z_{53} \text{X5P}^p &= flux_{Ru5} \text{Ru5P}^m && \text{(7c)} \\
Z_{34}\, \text{Ru5P}^p + Q_8 S^p_{3-7} &= (Q_4 + Q_7 + Z_{43}) \text{R5P}^p && \text{(7d)} \\
Z_{35}\, \text{Ru5P}^p + c(Q_8)^p + c(Q_6)^p &= (Q_5 + Q_7 + Z_{53}) \text{X5P}^p && \text{(7e)}
\end{aligned}$$

When focusing on eqns (7a) and (7c), normalizing these equations by the fluxes leaving the compartments allows for expressing the labeling on G6P and Ru5P as a function of the labeling on the other metabolites of the subsystem:

$$G^m = \frac{G_{input}}{flux_g} T^m_{G6P} + \frac{Z_{13}}{flux_g} F^m \tag{8}$$

$$\mathrm{Ru5P}^p = \frac{Z_3 T^p_5}{flux_{Ru5}} + \frac{Z_{43}\mathrm{R5P}^p}{flux_{Ru5}} + \frac{Z_{53}\mathrm{X5P}^m}{flux_{Ru5}} \tag{9}$$

In a next, crucial step, we substitute for $G^m$ and $\mathrm{Ru5P}^p$ with equations (8) and (9) in the remaining equations of the subsystem (7). After this step, they no longer depend on G6P and Ru5P while the tracer balances for F6P, R5P, and X5P are still valid. However, the modified equations contain elaborate fractions of the Z-fluxes, which still need to be simplified. For example, after some rearrangement eqn (8) inserted in (7b) results in:

$$Z_{12}\frac{G_{input}}{flux_g} T^m_{G6P} + F_{input} T^m_{F6P} + Q^m_{inp} = \left( Z_{13}\frac{Z_3}{flux_g} + Q_{loss} \right) F^m \tag{10}$$

Combining the first two terms on the left of eqn (10) gives us an input-derived flow, while the first term on the right refers to a flow from F6P to G6P that is followed by the first step of the oxidative PPP pathway. Thus we set:

$$\bar{Q}_1 = \frac{G_{input} Z_{12}}{Z_{12} + Z_3} + F_{input}; \quad \bar{Q}_3 = \frac{Z_3 Z_{13}}{Z_3 + Z_{12}}; \quad \bar{Q}_{12} = \frac{G_{input} Z_3}{flux_g} \tag{11}$$

Based on eqn (8), the transfer from G6P to the oxidative PPP via $Z_3$ is:

$$Z_3 G^m = \frac{G_{input} Z_3}{flux_g} T^m_{G6P} + \frac{Z_3 Z_{13}}{flux_g} F^m = \bar{Q}_{12} T^m_{G6P} + \bar{Q}_3 F^m \tag{12}$$

As a consistency check, we can derive

$$\bar{Q}_1 + \bar{Q}_{12} = \frac{G_{input} Z_{12}}{Z_{12} + Z_3} + F_{input} + \frac{G_{input} Z_3}{Z_{12} + Z_3} = G_{input} + F_{input} \tag{13}$$

$Q_{12}$ reflects that fraction of $G_{input}$ that is converted to Ru5P without reaching the F6P pool. Based on the preceding substitutions it follows that the hexose patterns are equal to the F6P patterns of the basic system.

We now focus on the pentose patterns. Replacing the Ru5P isotopomers $\mathrm{Ru5P}^p$ in eqn (7d) and (7e) with terms shown in eqn (9) while using the following substitutions

$$\frac{Z_{34}}{flux_{Ru5}} = p_R; \quad \frac{Z_{35}}{flux_{Ru5}} = p_X; \tag{14}$$

results in

$$Z_{34}\mathrm{Ru5P}^p - Z_{43}\mathrm{R5P}^p = p_R Z_3 T_5^p + p_R Z_{53}\mathrm{X5P}^p - p_X Z_{43}\mathrm{R5P}^p \tag{15}$$

for the exchange between Ru5P and R5P, and

$$Z_{35}\mathrm{Ru5P}^p - Z_{53}\mathrm{X5P}^p = p_X Z_3 T_5^p + p_X Z_{43}\mathrm{R5P}^p - p_R Z_{53}\mathrm{X5P}^p \tag{16}$$

for the exchange between Ru5P and X5P, respectively. With $p_R Z_{53} = BX$ and $p_X Z_{43} = BR$, one can define reduced balances for the R5P and X5P subsystem:

$$\begin{aligned} p_R Z_3 T_5^p + BX\,\mathrm{X5P}^p + Q_8 S_{3-7}^i &= (Q_4 + Q_7 + BR)\mathrm{R5P}^p \\ p_X Z_3 T_5^p + BR\,\mathrm{R5P}^p + c(Q_8)^p + c(Q_6)^p &= (Q_5 + Q_7 + BX)\mathrm{X5P}^p \end{aligned} \tag{17}$$

When summing up over all isotopomers, one obtains the flux balances for R5P and X5P:

$$\begin{aligned} p_R Z_3 + BX - BR + Q_8 &= Q_4 + Q_7 \\ p_X Z_3 - BX + BR + Q_8 + Q_6 &= Q_5 + Q_7 \end{aligned} \tag{18}$$

Thus, the balances of R5P and X5P require the three additional parameters $p_R$, $BR$, and $BX$, which define the exchange between the pentose pools and the NADPH oxidase-linked flow $Z_3$ that feeds into these pools. If necessary, they can be used to asses $Z_{53}$, and $Z_{43}$ as:

$$\mathrm{X5P} \xrightarrow{BX} \mathrm{R5P}: \; Z_{53} = \frac{BX}{p_R}; \qquad \mathrm{R5P} \xrightarrow{BR} \mathrm{X5P}: \; Z_{43} = \frac{BR}{p_X} \tag{19}$$

The resulting core system is shown in Figure 2. Given the flows and labeling patterns for the core system, one can assess the G6P and Ru5P patterns with eqn (7a) and (9), respectively. The flux $Z_3$ results from the sum over all isotopomers of equation (12) with $Z_3 = Q_3 + Q_{12}$. These equations serve as an interface to calculate values of the original system while using data of the core system.

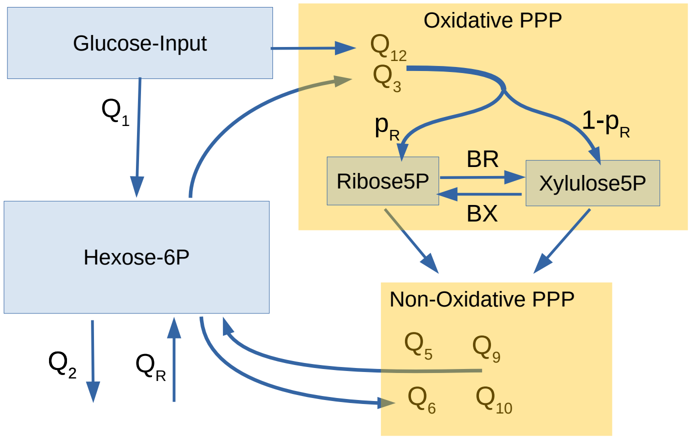


Figure 2: The condensed system. It affects the exchange between F6P and G6P and the flow through the NADPH oxidase step. The input is split into a 'direct' use by the NADPH oxidase ($Q_{12}$) and an uptake into the hexose pool $Q_1$.

#### 4.1 Intermediates in a reaction chain with reversible steps

A special situation arises for an unbranched reaction chain across 3 metabolites with a net transfer from starting to end point. The middle metabolite of the chain may be in a reversible exchange between the first and last metabolite with its labeling being somewhere in between. While the reversible exchange determines the label on the middle metabolite, it does not affect the net flow or the labeling on the start and end points. Therefore, it can be disregarded when formulating the network equations. In the case that measurements are available for the middle metabolite, its labeling pattern can be calculated from the values of the core system.

##### 4.1.1 6-Phosphogluconate (6PG)

6PG can be created from G6P. Britt et al. [12] have suggested that the subsequent NADPH oxidase step can be reversible, e.g. as a corresponding 'reverse' step starting from Ru5P. This 'reverse' flow, denoted with $k_1$, combines Ru5P and cytosolic $CO_2$. The transfer of the condensation product covering all isotopomers mass offset m that are derived from a condensation of labeled

Ru5P and cytosolic $^{13}CO_2$ is denoted as $c(k_1)^m = k_1[Ribulose \otimes {}^{13}CO_2]^m$. This results in the following mass balance for 6PG:

$$Z_3\,\mathrm{G6P}^m + c(k_1)^m = (k_1 + Z_3)\,\mathrm{6PG}^m$$

or

$$\mathrm{6PG}^m = \frac{Z_3\,\mathrm{G6P}^m + c(k_1)^m}{k_1 + Z_3} \tag{20}$$

Thus, the enrichment of the isotopomer $\mathrm{6PG}^m$ results from the sum of the enrichments on G6P and a backflow via the condensation of Ru5P with $CO_2$. Note that the labeling on G6P and Ru5P can be derived from the core metabolites with eqns (8) and (9). Two additional unknown parameters arise for the backflow $k_1$ and the cytosolic $^{13}CO_2$ labeling.

###### 4.1.2 Fructose-1,6-biphosphate (F-1,6-BP)

F-1,6-BP can be formed through phosphorylation of F6P and with a reversible condensation of GAP and DHAP.

Fluxes $Q_2$ and $Q_R$ apply to the entire reaction sequence from F6P to the two trioses. These steps are regulated between F6P and F-1,6-BP, while the reversible step from F-1,6-BP down to the trioses is superimposed with the exchange flux $k_{ex}$. Similarly to 4.1.1, we denote the flux of isotopomers with mass offset m resulting from the condensation of GAP and DHAP as $c(Q_R + k_{ex})^m = (Q_R + k_2)[GAP \otimes DHAP]^m$. We obtain the following mass balances for F-1,6-BP:

$$\text{Input} = \text{Output}$$

$$Q_2\mathrm{F6P}^m + c(Q_R + k_{ex})^m = (Q_R + Q_2 + k_{ex})(\text{F-1,6-B})^m$$

or

$$(\text{F-1,6-B})^m = \frac{Q_2\mathrm{F6P}^m + c(Q_R + k_{ex})^m}{Q_R + Q_2 + k_{ex}} \tag{21}$$

### 5 Constraints, based on flux balance equations

For the different pools of the system in Fig. 1, input-output balance equations can be defined. Only the net fluxes of reversible reactions are of relevance and denoted with the symbol $\Delta$ in the following. Specifically, we use $\Delta Q2 = Q_2 - Q_R$ for the net glycolytic rate, and $\Delta TAL$ for the transaldolase net flow $Q_9 - Q_{10}$. Further definitions include $\Delta TKT2 = Q_5 - Q_6$, $\Delta TKT1 = Q_7 - Q_8$, $\Delta GPI = Z_{12} - Z_{13}$ (glucose-6-phosphate-isomerase), $\Delta TPI = Q_{13} - Q_{14}$(triose-phosphate-isomerase), and $\Delta PEX = BR - BX$. With these net fluxes, the mass balances for the core system and the G6P compartment are

$$
\begin{aligned}
\text{G6P: } & Z_1 - F_{Input} \;=\; Z_3 + \Delta GPI \\
\text{F6P: } & F_{Input} + \Delta GPI + \Delta TAL + \Delta TKT2 \;=\; \Delta Q2 \\
\text{R5P: } & p_R Z_3 \;=\; Q_4 + \Delta PEX + \Delta TKT1 \\
\text{X5P: } & p_X Z_3 \;=\; \Delta TKT2 + \Delta TKT1 - \Delta PEX \\
\text{S7P: } & S_{inp} + \Delta TKT1 \;=\; \Delta TAL \\
\text{E4P: } & \Delta TAL \;=\; \Delta TKT2 \\
\text{DHAP: } & \Delta Q2 + T_{inp} \;=\; \Delta TPI \\
\text{GAP: } & \Delta Q2 + \Delta TPI + \Delta TKT2 + \Delta TKT1 \;=\; Q_{11} + \Delta TAL
\end{aligned} \tag{22}
$$

These mass balances can be formally expressed as a matrix

$$0 = [\mathbf{M}_0] \times \mathbf{fluxes}$$

and this equation can then be split into two parts:

$$0 = [\mathbf{M}_a] \times \mathbf{fluxes_a} + [\mathbf{M}_b] \times \mathbf{fluxes_b}$$

For the next step we require the separation of matrix $[\mathbf{M}_0]$ to be performed in a way so that matrix $[\mathbf{M}_a]$ is square and invertible. This allows the following steps:

$$
\begin{aligned}
-[\mathbf{M}_a] \times \mathbf{fluxes_a} &= [\mathbf{M}_b] \times \mathbf{fluxes_b} \\
\mathbf{fluxes_a} &= -[\mathbf{M}_a]^{-1} [\mathbf{M}_b] \times \mathbf{fluxes_b} \\
\mathbf{fluxes_a} &= [\mathbf{M}_c] \times \mathbf{fluxes_b} \quad \text{with } [\mathbf{M}_c] = -[\mathbf{M}_a]^{-1} [\mathbf{M}_b]
\end{aligned} \tag{23}
$$

With these rearrangements, eqn (23) allows for defining 8 dependent fluxes ( $\mathbf{fluxes_a}$) and 7 independent fluxes $\mathbf{fluxes_b}$. $[\mathbf{M}_c]$ serves as a *Dependency* *matrix*, resulting in the following corresponding dependency system:

$$\begin{pmatrix} Z_3 \\ Q_4 \\ Q_{11} \\ \Delta GPI \\ \Delta TKT2 \\ \Delta TKT1 \\ \Delta TPI \\ \Delta PEX \end{pmatrix} = \begin{pmatrix} 1 & -1 & 2 & 0 & 0 & 0 & 0 \\ 1 & -1 & -1 & 1 & 0 & 2 & 0 \\ 0 & 2 & 1 & 0 & 1 & -1 & 0 \\ 0 & 1 & -2 & 0 & 0 & 0 & -1 \\ 0 & 0 & 1 & 0 & 0 & 0 & 0 \\ 0 & 0 & 1 & 0 & 0 & -1 & 0 \\ 0 & 1 & 0 & 0 & 1 & 0 & 0 \\ -pX & pX & 2pR & 0 & 0 & -1 & 0 \end{pmatrix} \times \begin{pmatrix} Z_1 \\ \Delta Q2 \\ \Delta TAL \\ P_{Input} \\ T_{input} \\ S_{input} \\ F_{Input} \end{pmatrix} \quad (24)$$

#### 5.1 Application of flux constraints to extreme conditions

We assume that the minimal value of $\Delta Q2$ is -20, which would indicate the gluconeogenetic flux $Q_R$ exceeding the glycolytic flux $Q_2$ by 20. We further set $Z_1 = 100$. In the dependency system, only the dependent fluxes $Z_3, Q_4$ and $Q_{11}$ are subject to the restriction that they do not become negative. These restrictions play a role in potential extreme values and are calculated with the first 3 rows of the dependency system. The following discussion refers to these lines. In our scenario, the input flows appear only in combination, namely:

$$Pent_{input} = P_{Input} + 2S_{Input}; \quad Triose_{Input} = T_{Input} - S_{Input} \quad (25)$$

##### 5.1.1 Scenario: No R5P loss from the network

$Q_4 = 0$ implies a complete utilization of the entire hexose input either via glycolysis or NADPH oxidation. We set $Q_4 = 0$ for the second row of eqn (24):

$$\begin{aligned} Q_4 = 0 =& Z_1 - \Delta Q2 - \Delta TAL + Pent_{Input} \\ \text{or:} \quad \Delta TAL =& Z_1 - \Delta Q2 + Pent_{Input} \end{aligned} \quad (26)$$

When replacing $\Delta TAL$ with eqn (26), the first and third row of eqn (24) result in:

$$\begin{aligned} Z_3 =& Z_1 - \Delta Q2 + 2(Z_1 - \Delta Q2) + Pent_{Input} \\ =& 3(Z_1 - \Delta Q2) + Pent_{Input} \quad (27a) \\ Q_{11} =& 2\Delta Q2 + \Delta TAL + Triose_{Input} \\ =& 2\Delta Q2 + (Z_1 - \Delta Q2) + Pent_{Input} + Triose_{Input} \\ =& Z_1 + \Delta Q2 + Pent_{Input} + Triose_{Input} \quad (27b) \end{aligned}$$

If we now disregard the input values, we get the following values for this extreme scenario: For $\Delta Q2 = 0$ the maximum TAL value would be 100, the maximum $Z_3$ value 300, and the minimum $Q_{11}$ value 100. These limits can be shifted by $\Delta Q2$.

###### 5.1.2 Scenario: Ribose formation exclusively through the non oxidative PPP

In this scenario, the complete input $Z_1$ must be converted to R5P through the non-oxidative PPP and leave the network. Accordingly, if we set $Z_3 = 0$ and $Q_{11} = 0$, then

$$\begin{aligned} Q_{11} = 0 =& 2\Delta Q2 + \Delta TAL + Triose_{Input} \\ \text{or: } \Delta Q2 =& - \Delta TAL/2 - Triose_{Input} \quad (28) \end{aligned}$$

When using this expression for $\Delta Q2$ and setting $Z_3 = 0$, one obtains

$$\begin{aligned} Z_3 = 0 =& Z_1 - \Delta Q2 + 2\Delta TAL \\ \text{or: } 0 =& Z_1 + \Delta TAL/2 + 2\Delta TAL - Triose_{Input} \\ \text{or:} \quad \Delta TAL =& - Z_1/2.5 + Triose_{Input}/2.5 = -40 + Triose_{Input}/2.5 \quad (29) \end{aligned}$$

from the first row of eqn (24). Similarly, we get the following equations for the Ribose formation via $Q_4$:

$$Q_4 = Z_1 - \Delta Q2 - \Delta TAL + Pent_{Input} = Z_1 + \Delta TAL/2 - \Delta TAL + Pent_{Input}$$

$$Q_4 = Z_1 - \Delta TAL/2 + Pent_{Input} \; = \; Z_1 + Z_1/5 = \frac{6}{5}Z_1 + Pent_{Input} \quad (30)$$

From eqn (30), it can be deduced that 6 mol of R5P can be maximally produced by 5 mol of hexose. Furthermore, the minimal flux of $\Delta TAL$ is

-100/2.5 or -40. Under these conditions $Q_4$ would be 120, while 20 of it (from eqn (28)) would be derived from the triose formation by $\Delta Q2$ or the first steps of glycolysis. So far, the derivation indicates that $\Delta TAL$ ranges from -40 to 100. These extremes include complete hexose utilization for R5P production and extend to complete oxidation of the upper half of hexose and loss of the lower half from the PPP as triose.

### 6 Sampling for the core system

In a Bayesian approach [13], prior ranges or initial distributions can be specified for individual fluxes. These priors should be as 'non-informative' as possible and the allowed parameter range should be fully explored/covered by the prior distributions. However, there is one restriction to take into consideration: fluxes derived from other parameters should never become negative, as negative fluxes can lead to a program termination. In the previous section, we derived upper and lower bounds for selected fluxes that fulfill these constraints. In the following, we now develop an approach to define admissible flux or parameter ranges.

#### 6.1 Parameter and flux sampling

With the lower limit $\Delta Q2 = -20$, the first line of the dependency equation (24) results from:

$$Z_3 = Z_1 + 20 + 2\Delta TAL \; = \; 120 + 2\Delta TAL \tag{31}$$

$Z_3$ becomes smaller when $\Delta Q2$ becomes larger than the limit value of -20. Thus, a tolerable $Z_3$ is always below on the limit line defined by eqn (31). The flux $Q_4$, which we want to limit from becoming negative, is defined by:

$$Q_4 = Z_1 - \Delta Q2 - \Delta TAL + Pent_{input} \tag{32}$$

We resolved the first line of the equation system (24) for $\Delta Q2$

$$\Delta Q2 = Z_1 - Z_3 - 2\Delta TAL \tag{33}$$

and used this expression to replace $\Delta Q2$ in eqn (32):

$$\begin{aligned} Q_4 =& Z_1 - (Z_1 - Z_3 + 2\Delta TAL) - \Delta TAL + Pent_{input} \\ =& Z_3 - 3\Delta TAL + Pent_{input} \end{aligned} \tag{34}$$

This results in the limit line $Z_3 - 3\Delta TAL + Pent_{input} = 0$. When rearranging this formula to

$$Z_3 = 3\Delta TAL - Pent_{input} \tag{35}$$

we gain another limit line that $Z_3$ needs to exceed. A similar strategy can be utilized for $Q_{11}$.

$$\begin{aligned} Q_{11} =& 2\Delta Q2 + \Delta TAL + Triose_{inp} \quad \text{using eqn (33) gives} \\ =& 2(Z_1 - Z_3 + 2 * \Delta TAL) + \Delta TAL + Triose_{inp} \\ =& 2Z_1 - 2Z_3 + 5\Delta TAL + Triose_{inp} \end{aligned} \tag{36}$$

If we set $Q_{11}$ to zero, it results in a third and final limit line, which $Z_3$ must be below.

$$Z_3 = Z_1 + \frac{5}{2}\Delta TAL + \frac{1}{2}Triose_{inp} \tag{37}$$

Figure 3 demonstrates the resulting feasible range for sampling $Z_3$ and $\Delta TAL$.

Sampling is initiated by drawing random non-negative values for $T_{input}$, $S_{input}$ and $P_{input}$. With these input values, the feasible range for $Z_3$ and $\Delta TAL$ can be defined with eqns (31), (35) and (37). We first sampled from the $\Delta TAL$ range, represented by the blue, dashed horizontal line in Fig. 3. By indicating the sampled $\Delta TAL$ by a horizontal arrow in the graphic, we can define a vertical dashed line through the yellow segment reflecting the sampling range for corresponding $Z_3$ values. An example for a potential $Z_3$ value is given by the vertical arrow. After sampling of $\Delta TAL$ and $Z_3$, $\Delta Q2$ can be determined using eqn (33). Then, the five different $\Delta$-net fluxes can be calculated with eqn (24). These net fluxes must be converted into forward and reverse fluxes, such as:

$$Q_6 = -\Delta TKT2 + Q_5; \quad Q_8 = -\Delta TKT1 + Q_7; \tag{38}$$

Therefore, 5 more 'independent' fluxes ( $Q_5$, $Q_7$, $Q_{13}$, $Z_{13}$ and $BR$) must be collected so that both the dependent ( $Q_6$, $Q_8$, $Z_{12}$, $Q_{14}$ and $BX$ ) and independent fluxes are always greater than or equal to zero. Hence, we split the 'independent fluxes/parameters' into

$$\begin{aligned} Q_6 &= -\Delta TKT2 + Q_{5,min} + Q_{5,offset} \\ Q_8 &= -\Delta TKT1 + Q_{7.min} + Q_{7,offset} \end{aligned} \tag{39}$$

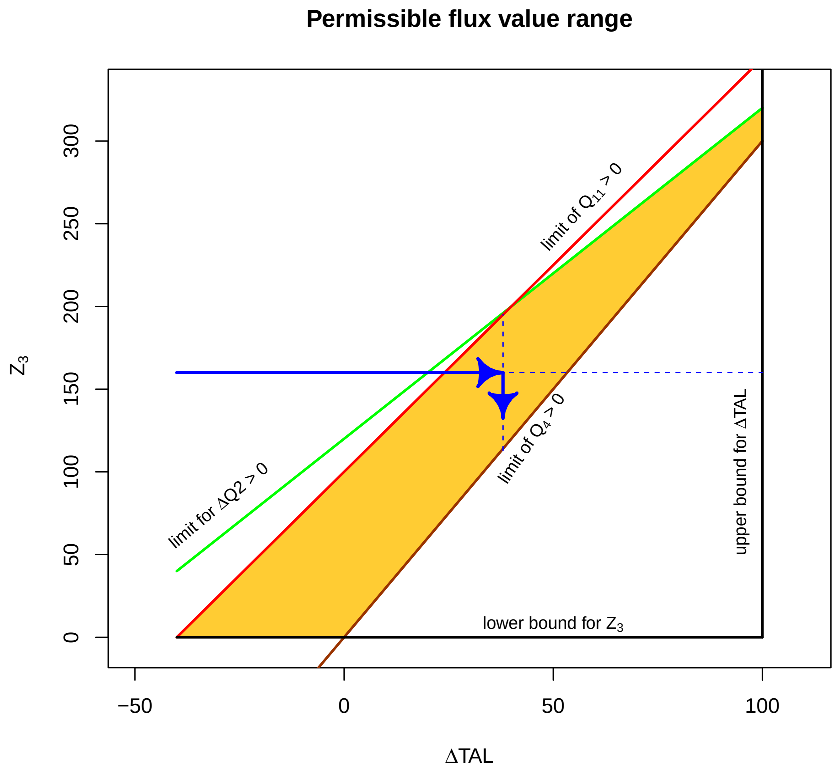


Figure 3: Permissible range (orange polygon) for the parameters $Z_3$ and $\Delta TAL$ in the case that the input values are zero.

where the offset values are always $\geq 0$. If the $\Delta$-values are positive (e.g. $\Delta TKT2$), we set $Q_{5,min} = \Delta TKT2$, else $Q_{5,min} = 0$ . With this strategy, dependent and independent fluxes are always $\geq 0$.

### 7 Testing for correct enrichment calculation

The EMU approach requires individual calculation steps for each level. For own implementations, a programming error can creep in on each level. Therefore, we use a test to verify whether patterns at higher levels are compatible with those from lower levels. For this test we require the following quantities: a) The abundance on isolated carbons, independent of the abundance on adjacent carbons. We define this quantity as $\hat{c}_i(metabolite)$, i.e $\hat{c}_i(G6P)$, where the index i refers to the carbon position. b) The $^{13}C$ content of a fragment ion. It is denoted and calculated as:

$$\bar{C}_{a-b} = 1/n \sum_{1}^{n} i * p_{a-b}^{i} \tag{40}$$

where $p_{a-b}^{i}$ is the abundance of isotopomers with mass offset i across the fragment ranging from carbon a to carbon b and n refers to the total number of considered carbons. As outlined by Lima et al. [14], the $^{13}$C -content of a fragment can be related to the isolated carbon labeling of a fragment:

$$\sum_{i=1}^{n} \hat{c}_i = n\bar{C}_{1-n} \tag{41}$$

The isolated carbon labeling of a given metabolite is obtained from EMU level 1. Each subsequent EMU level adds one additional carbon to the distribution. Thus, for each calculated distribution over a longer segment of the carbon skeleton, there is the corresponding set of level-1 enrichments and a test based on equation (41) can be performed. While these tests might only capture whether an EMU setup is consistent across different levels, implementing them is warranted as these inconsistencies pose a critical obstacle during implementation.

### References


[1] Antoniewicz MR, Kelleher JK, Stephanopoulos G. Elementary Metabolic Units (EMU): a novel framework for modeling isotopic distributions. Metab Eng. 2007;9(1):68–86.

[2] Wiechert W, Wurzel M. Metabolic isotopomer labeling systems: Part I: global dynamic behavior. Mathematical Biosciences. 2001;169(2):173 – 205.

[3] Carpenter B, Gelman A, Hoffman MD, Lee D, Goodrich B, Betancourt M, et al. Stan: A probabilistic programming language. Journal of statistical software. 2017;76(1).

[4] Stan Development Team. The Stan Core Library; 2023. Version 2.32.0. Available from: `https://mc-stan.org/users/documentation/`.

[5] Stan Development Team. RStan: the R interface to Stan; 2023. R package version 2.21.8. Available from: https://mc-stan.org/.

[6] Stan Development Team. RStan: the R interface to Stan, Version 2.17.3; 2018. Available from: http://mc-stan.org/rstan.html.

[7] Metropolis N, Rosenbluth AW, Rosenbluth MN, Teller AH, Teller E. Equation of State Calculations by Fast Computing Machines. J Chem Phys. 1953;21:1087–1092.

[8] Hoffman MD, Gelman A. The No-U-Turn Sampler: Adaptively Setting Path Lengths in Hamiltonian Monte Carlo. Journal of Machine Learning Research. 2014;15:1593–1623.

[9] Stan Development Team; 2020.

[10] Bishop CM. Pattern Recognition and Machine Learning (Information Science and Statistics). 1st ed. Springer; 2007.

[11] Katz J, Rognstad R. The labeling of pentose phosphate from glucose-14C and estimation of the rates of transaldolase, transketolase, the contribution of the pentose cycle, and ribose phosphate synthesis. Biochemistry. 1967 Jul;6(7):2227–2247.

[12] Britt EC, Lika J, Giese MA, Schoen TJ, Seim GL, Huang Z, et al. Switching to the cyclic pentose phosphate pathway powers the oxidative burst in activated neutrophils. Nature Metabolism. 2022 Mar;4(3):389–403. Available from: https://doi.org/10.1038/s42255-022-00550-8.

[13] Gelman A, Carlin JB, Stern HS, Dunson DB, Vehtari A, Rubin DB. Bayesian Data Analysis, Third Edition. Chapman and Hall, New York; 2013.

[14] Lima VF, Erban A, Daubermann AG, Freire FBS, Porto NP, Cândido-Sobrinho SA, et al. Establishment of a GC-MS-based (13) C-positional isotopomer approach suitable for investigating metabolic fluxes in plant primary metabolism. Plant J. 2021 Sep;108(4):1213–1233.
